# Rapid liquid biopsy assessment through gene profiling from the kidney biopsy transport medium: a technical validation and a proof-of-concept pilot study

**DOI:** 10.1101/2024.07.28.604919

**Authors:** Ziyang Li, Marij J.P. Welters, Hailiang Mei, Alexis Varin, Baptiste Lamarthée, Aiko P.J. de Vries, Hans J. Baelde, Jesper Kers

**Affiliations:** Department of Pathology, and Leiden Transplant Center, Leiden University Medical Center, Leiden, The Netherlands; Department of Medical Oncology, Oncode Institute, Leiden University Medical Center, Leiden, The Netherlands; Sequencing Analysis Support Core, Leiden University Medical Center, Leiden, The Netherlands; Université Marie et Louis Pasteur, INSERM UMR Right, Établissement Français du Sang, Besançon, France; Department of Internal Medicine, Division of Nephrology, and Leiden Transplant Center, Leiden University Medical Center, Leiden, The Netherlands; Department of Pathology, Amsterdam UMC, Amsterdam, The Netherlands; Department of Pathology, Erasmus MC, Rotterdam, The Netherlands

**Author notes:** <u>Corresponding author:</u> Ziyang Li, Department of Pathology, Leiden University Medical Center, Albinusdreef 2, 2333 ZA, Leiden, The Netherlands.

**Keywords:** biopsy transport medium, gene profiling, fast molecular diagnosis, kidney transplantation

## Abstract

Rapid diagnosis is pivotal in kidney disease for timely and precision therapy. Conventional microscopic and molecular assessments from biopsy tissues rely on extra sample processing, making same-day diagnosis impractical. Therefore, we introduce the biopsy transport medium (BTM), a byproduct of the biopsy tissue storage process that could serve as a source of biomarkers, accelerating the assessment workflow. Biopsies from tumor-free nephrectomy tissues were used to create mimicked BTM, allowing optimization of RNA extraction procedure. RNA yield and integrity were then systematically evaluated prior to downstream analyses. Subsequently, gene expression analysis was performed through multiple techniques: qPCR, RNA sequencing, and NanoString nCounter system. The results showed that storage time (the duration a biopsy is stored in BTM), ranging from 0.5 to 24 hours, did not significantly affect RNA quality and yield. The transcriptomic signals detected in biopsy tissues are largely recapitulated in the corresponding BTM samples. The differential gene expression analysis based on BTM identified rejection-associated profiles, which are aligned with Banff lesion scores. This study confirms BTM’s ability to provide transcriptomic information relevant to the state of the kidney and supports BTM’s potential for same-day molecular diagnosis, especially with tailored qPCR panels for rapid, targeted analysis.

## 1 INTRODUCTION

Rapid diagnosis is imperative for effectively managing kidney diseases, particularly in cases of acute kidney injury (AKI), which poses a significant threat to future organ function, akin to the critical nature of myocardial infarction and cerebral stroke scenarios (’time equals tissue’). AKI cases with high inflammatory activity, such as acute rejection in transplants, demand immediate intervention (e.g., plasma exchange or immunosuppressive therapy) to prevent irreversible damage, whereas treatment delays risk tissue loss, fibrosis, and progressive functional decline.

The current gold standard for assessing disease status involves microscopic examination after formalin fixation and tissue processing, which typically takes at least 24 hours for an initial assessment. Although frozen section pathology in principle enables quick results, its resolution and accuracy are challenged, especially when handling complex specimens. Traditional histology can be further delayed by variability and differing interpretations among pathologists. Emerging molecular technologies offer potential advancements, such as the Molecular Microscope Diagnostic System (MMDx) and the NanoString Banff Human Organ Transplant (B-HOT) targeted gene profiling system. Nevertheless, MMDx requires additional biopsy cores, and B-HOT profiling necessitates further sample processing of formalin-fixation, paraffin-embedding (FFPE) material, limiting their accessibility in clinical practice^1,2^. Typically, molecular profiling is only pursued if histology results remain equivocal.

In response to these limitations, we are currently exploring alternative methods to overcome these barriers by investigating a novel and freely accessible liquid biopsy. At the Leiden University Medical Center (LUMC), fresh kidney biopsies are promptly transported to the pathology ward floating in phosphate-buffered saline (PBS). The pathology department processes the biopsy, and the biopsy transport medium (BTM) is typically discarded. The BTM, however, potentially contains a diverse array of tissue-derived molecules, vesicles, and live cells, providing a broad range of biological information for diagnostic purposes and potentially functional tests.

In this study, our objectives were: 1) to validate the technical feasibility of utilizing the BTM as a source of molecular information, and 2) to present a proof-of-concept study demonstrating that gene signature profiling from the BTM could provide actionable information to classify rejection (subtypes) versus non-rejection in the commonly encountered clinical scenario of delayed graft function (DGF).

## 2 MATERIALS and METHODS

### 2.1 Preliminary experiments

We prospectively collected and processed BTM derived from real kidney biopsies from the pathology department in LUMC for later selection. All the materials were obtained, completely anonymized (i.e. no pseudonymization code was put in place and no clinical data were retrieved from the electronic health records), and handled according to institutional guidelines, Good Research Practice, in accordance with Dutch national ethics guidelines according to (the Code of Conduct for the Proper Secondary Use of Human Tissue). BTM was centrifuged at 300g for 5 minutes to collect the sediment. The pellet was resuspended in 1 mL of Red Blood Cell Lysis Buffer (Roche Diagnostics, Mannheim, Germany) and gently inverted periodically for 5–10 minutes at room temperature. The samples were then centrifuged at 500g for 5 minutes at room temperature. Following centrifugation, the supernatant was removed, leaving the white cell pellet, which was subsequently used for cytospin-based morphological assessment and flow cytometry analysis of real BTM.

### 2.2 Mimic BTM Samples and processing

The kidney biopsy procedure at LUMC typically involves two routine needle passes, utilizing a 16G needle. Two biopsies are then acquired, each measuring 15-20 mm in length and approximately 1 mm in diameter. To standardize the condition of the BTM, we created the kidney biopsy by harvesting tumor-free tissues from a tumor nephrectomy. To mimic the standard operating procedure (SOP) in our hospital, tissues were cut into slices of 1.5×1.5×20 mm, closely matching the dimensions of standard kidney biopsy cores, and two pieces of tissue were then placed in 15 mL PBS. In addition, two larger pieces of the adjacent tissue from the same nephrectomy were processed either as FFPE or fresh-frozen samples, the latter were stored at -80 □. The BTM without biopsy tissues was centrifuged at 1500g or 3000g to obtain the cell sediments, and the resulting cell sediments were lysed with 500 μl TRIzol™ Reagent (Ambion, Foster City, CA, USA) or 350 μl buffered RLT (included in RNeasy Micro Kit, QIAGEN, Hilden, Germany) + 3.5 μl 2-Mercaptoethanol (Merck Schuchardt OHG, Hohenbrunn, Germany) and homogenized. For fresh biopsy tissues, adjacent frozen tissues, and adjacent FFPE tissues, RNA isolation was performed using the TRIzol method. This choice was made due to the relatively large amount of starting material, while the RNeasy Kit was specifically designed for small sample volumes.

### 2.3 Optimization of RNA extraction protocols: a procedural comparison using mimicked BTM

To evaluate the effect of storage time (the duration a biopsy is floating in the BTM during transport/storage) on RNA quantity and integrity, we set up five groups with varying storage times, 0.5 hours, 1 hour, 2 hours, 4 hours, and 24 hours. To mimic clinical practice, samples were kept at room temperature for the first 0.5 hours and then transferred to a 4 °C refrigerator. At designated time intervals, biopsies were removed from the BTM, after which the residual material was centrifuged. To assess the effect of centrifugation speed, we set up two groups with centrifugation speeds of 1500g and 3000g for 0.5 hours, with the speed selection guided by reported protocols for RNA isolation from urine^3,4^. Plus, due to the relatively low RNA yield from BTM, we conducted both the TRIzol method and the RNeasy Micro Kit to compare extraction efficiencies. Before this, to optimize the aforementioned methods, we set up four isolation protocols: RNeasy without DNase treatment, RNeasy with DNase treatment (DNase treatment is an optional procedure in the manufacturer’s instructions), TRIzol method alone, and TRIzol method followed by RNeasy clean-up (QIAGEN, Hilden, Germany). An additional experiment was conducted to assess whether ultracentrifugation at 20,000g influences gene expression analysis. Hematoxylin and eosin (H&E) staining was performed on the corresponding tumor-free kidney FFPE tissues to evaluate tissue morphology.

### 2.4 PCR analysis of gene expression in mimicked BTM

The RNA with fixed volume was synthesized into cDNA with AMV reverse transcriptase (Roche, Basel, Switzerland) using random hexamer primers. qPCR analysis was performed utilizing IQTMSYBR Green Supermix (Bio-Rad, Hercules, CA, USA) in conjunction with the Bio-Rad CFX96 Touch Real-Time PCR Detection System and the Bio-Rad CFX Maestro 2.2 (5.2.008.0222) software. Since qPCR reflects transcript abundance, the RNA quantity was indirectly estimated through the Ct value of the housekeeping gene hypoxanthine phosphoribosyltransferase 1 (*HPRT1*). Additionally, because successful qPCR amplification highly depends on RNA integrity, the detectability of *HPRT1* also serves as an indirect indicator of RNA quality and its suitability for downstream PCR-based applications. To elucidate the distribution of cells released into the BTM, qPCR was used to detect the expression of eight cell-type-specific genes (*CD20, CD3E, CD56, CD68, CD138, PECAM1, NPHS1*, and *LRP2*) in all mimicked BTM. These genes serve as biomarkers for B cells, T cells, NK cells, macrophages, plasma cells, endothelial cells, podocytes, and tubular epithelial cells, respectively (Table 1). The expression levels of the target genes in each sample were normalized by *HPRT1* and calculated using the ΔΔCt method.

**Table 1.**
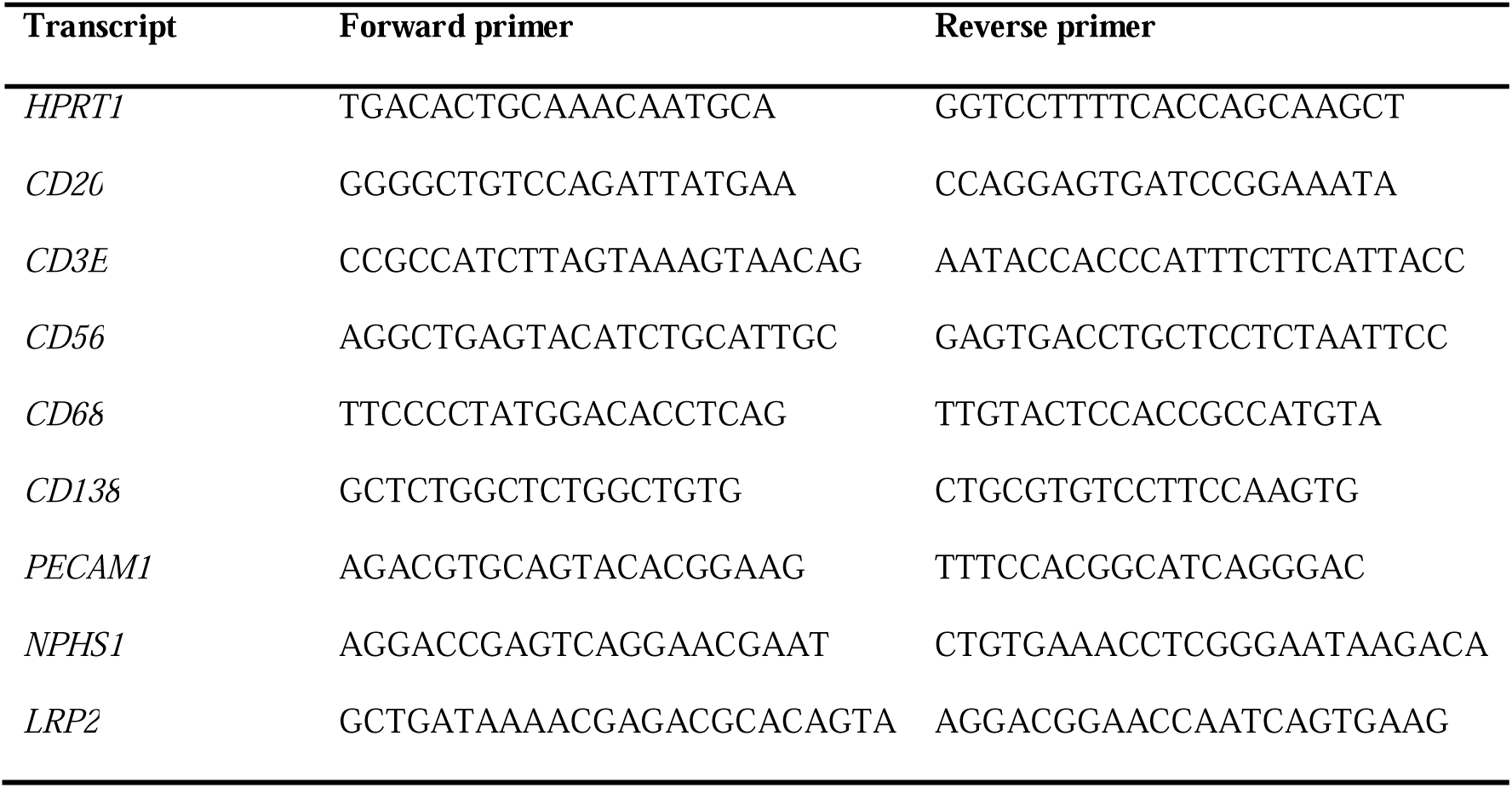
Primers.

### 2.5 Bulk RNA Sequencing (RNA-Seq) and NanoString B-HOT Panel Gene Expression Analysis

Bulk RNA-Seq was performed on mimicked BTM, fresh kidney biopsies, and adjacent frozen tissues. In parallel, gene expression profiling was conducted on mimicked BTM, corresponding frozen and FFPE tissues, and real BTM using the NanoString B-HOT panel (770 genes)^5^.

### 2.6 Statistical analysis

Group comparisons were evaluated using one-way ANOVA and Independent-Samples t-tests. Reliability and agreement were assessed using intraclass correlation coefficients (ICC) and Bland–Altman analysis. Differential gene expression (DGE) analysis was conducted, and cell fractions were imputed using CIBERSORTx. Deconvolution was performed with a signature matrix derived from single-cell RNA sequencing of 16 kidney allograft biopsies, enabling the discrimination of 24 parenchymal and immune cell phenotypes^6^.

A full description of the experimental procedures is available in Supplementary Methods.

## 3 RESULTS

### 3.1 Presence of live cells in the real BTM

The real BTM samples were collected without prior knowledge of the pathology results. Cytospin-prepared Wright–Giemsa staining of BTM samples showed well-preserved cell morphology. Immune cells and epithelial cells can be observed (Figure 1a, Supplementary Figure 1). H&E staining of the corresponding biopsy tissue demonstrated mild tissue damage (Figure 1b), corroborating the cytospin findings. Flow cytometry of an alternative BTM sample showed 45% *CD45*LJ cells and 7.6% *LRP2*LJ cells among live, debris-excluded populations (Supplementary Figure 2).

**Figure 1.**
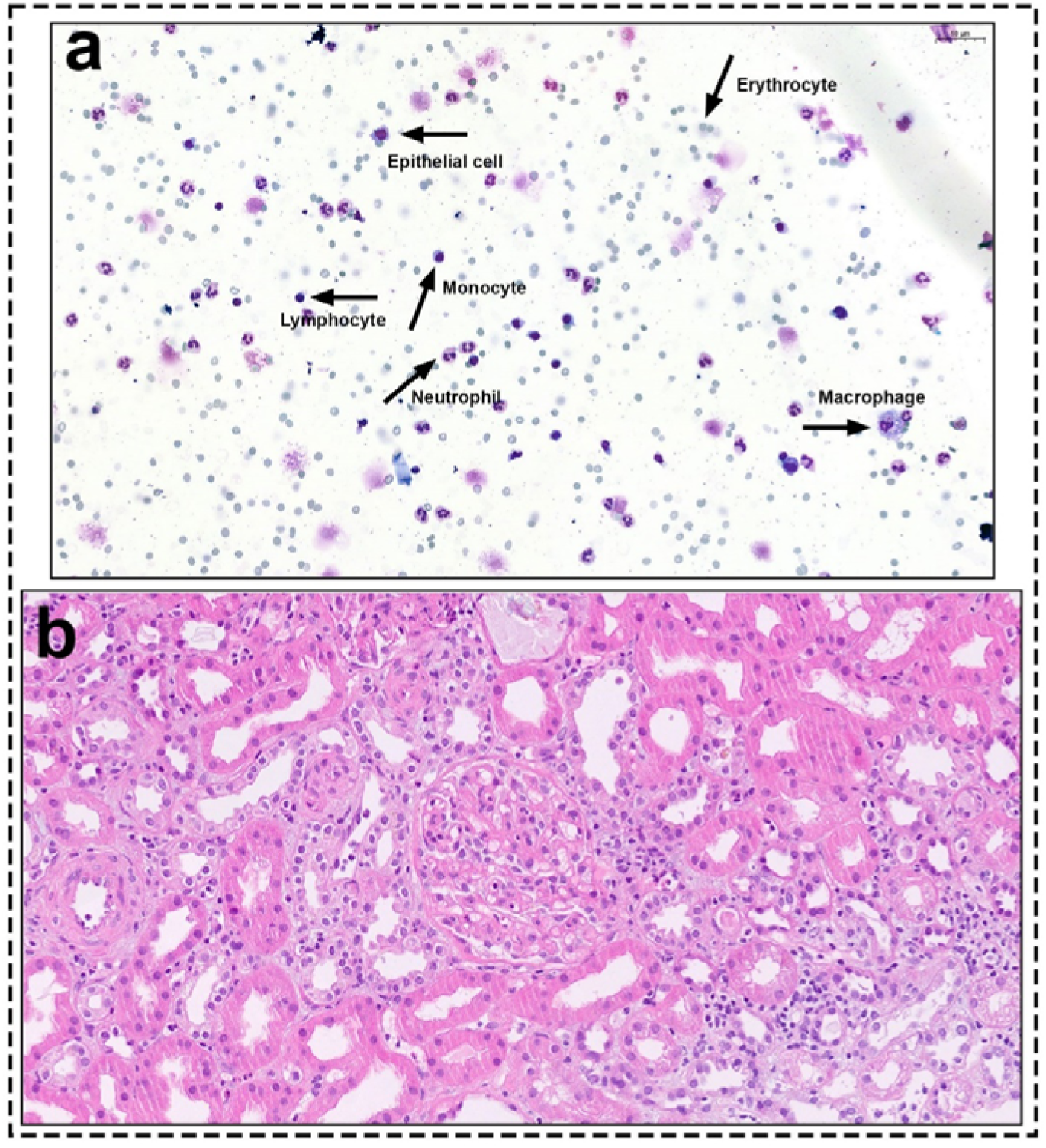
Cytospin-prepared Wright–Giemsa staining and matched FFPE tissue (H&E). (a) Wright–Giemsa staining reveals predominant cell types, including erythrocytes, monocytes, lymphocytes, neutrophils, macrophages, and, to a lesser extent, epithelial cells; (b) H&E staining demonstrates mild tissue damage in the form of focal interstitial inflammation and local tubular damage (epithelial vacuolization and simplification).

### 3.2 RNA yield of mimicked BTM

Tumor-free tissues were collected from 10 nephrectomies, and a total of 120 samples were made into mimicked BTM (12 identical samples were prepared from a single kidney). (Supplementary Figure 3). The mean RNA yield was 702 ± 613 ng across all samples. Storage time had no significant effect on yield (P = 0.92 in RNeasy-Kit group and P = 0.98 in TRIzol group, Figure 2a).

**Figure 2.**
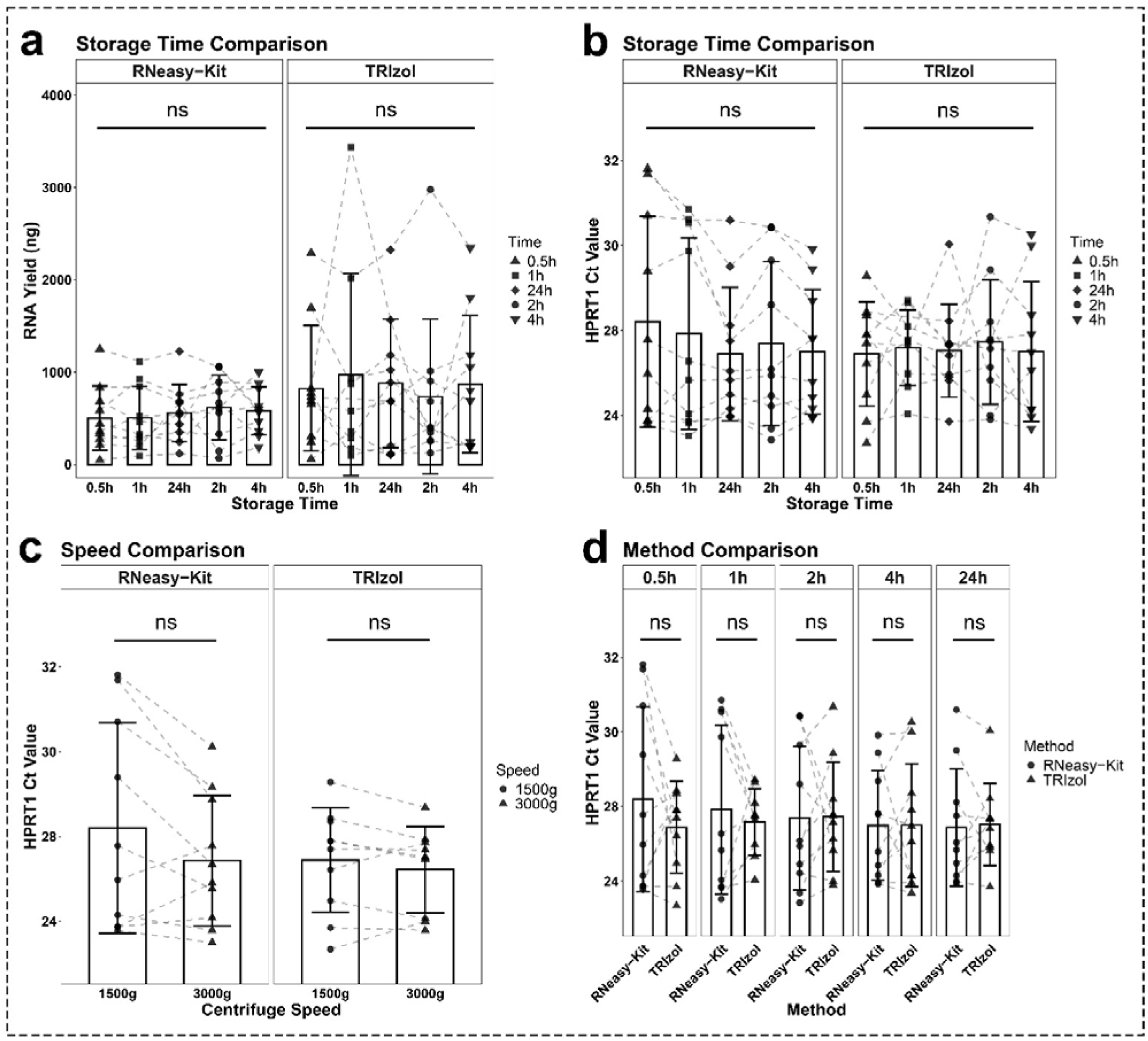
The procedure comparison on RNA yield and *HPRT1* Ct value. Dots denote individual samples. Lines connect paired samples within each group to illustrate within-group changes in RNA yields (a) or *HPRT1* Ct value (b, c, and d). Each line belonged to the samples from an individual kidney. There are no significant differences between subgroups within each group.

### 3.3 Procedural Effects on RNA Quantified by qPCR

RNA integrity was evaluated from 24 mimicked BTM samples divided into four isolation protocols (Supplementary Figure 4). Mean RINs were 5.66 ± 2.57 (TRIzol alone) vs. 7.43 ± 0.16 (TRIzol + Clean up, *P* = 0.20) and 5.18 ± 1.34 (RNeasy–DNase) vs. 7.55 ± 0.54 (RNeasy+DNase, *P* = 0.006). TRIzol + clean up, and RNeasy+DNase were selected for subsequent experiments. *HPRT1* was detected in a total of 120 mimicked BTM samples derived from 10 kidneys (mean Ct = 27.1 ± 2.12). *HPRT1* expression was consistent across storage times (0.5–24 h, P = 0.92 in RNeasy-Kit group and P = 0.98 in TRIzol group, Figure 2b), centrifugation speeds (1500 g vs. 3000 g, P = 0.42 in RNeasy-Kit group and P = 0.67 in TRIzol group, Figure 2c), and isolation methods (RNeasy vs. TRIzol, P = 0.40, 0.83, 0.96, 0.99, and 0.90 in different storage time groups, Figure 2d). Thus, these factors did not significantly affect RNA yield. Similarly, comparison of 1500 g vs. 20,000 g using three samples and four housekeeping genes showed no significant differences (P = 0.58, 0.71, 0.40, 0.24, Supplementary Figure 5). B-HOT panel analysis showed strong correlation across five storage time points (Supplementary Figure 6), indicating minimal effects of storage within 24 hours. The profiling of 770 genes further validated the qPCR findings.

### 3.4 Cell-type-specific marker expression measurement on mimicked BTM by qPCR

Our selected biomarkers encompass immune cells and parenchymal cells. While all markers are detectable in the BTM, there is a distinct cell distribution compared to corresponding frozen tissues (Figure 3). Normalized expression levels of immune cell-related biomarkers were all higher in BTM compared to in the corresponding frozen tissues, particularly *CD68* (4.25-fold increase, P = 0.004). Conversely, *NPHS1* expression (podocytes) is significantly lower in BTM compared to frozen tissues (5.63-fold decrease, P < 0.001). The expression of *PECAM1* (endothelial cells) and *LRP2* (proximal tubular epithelial cells) did not show significant differences between BTM and frozen tissue. FFPE sections of the corresponding 10 kidneys showed focal inflammation with immune cell infiltration (Supplementary Figure 7), consistent with the immune enrichment observed in mimicked BTM.

**Figure 3.**
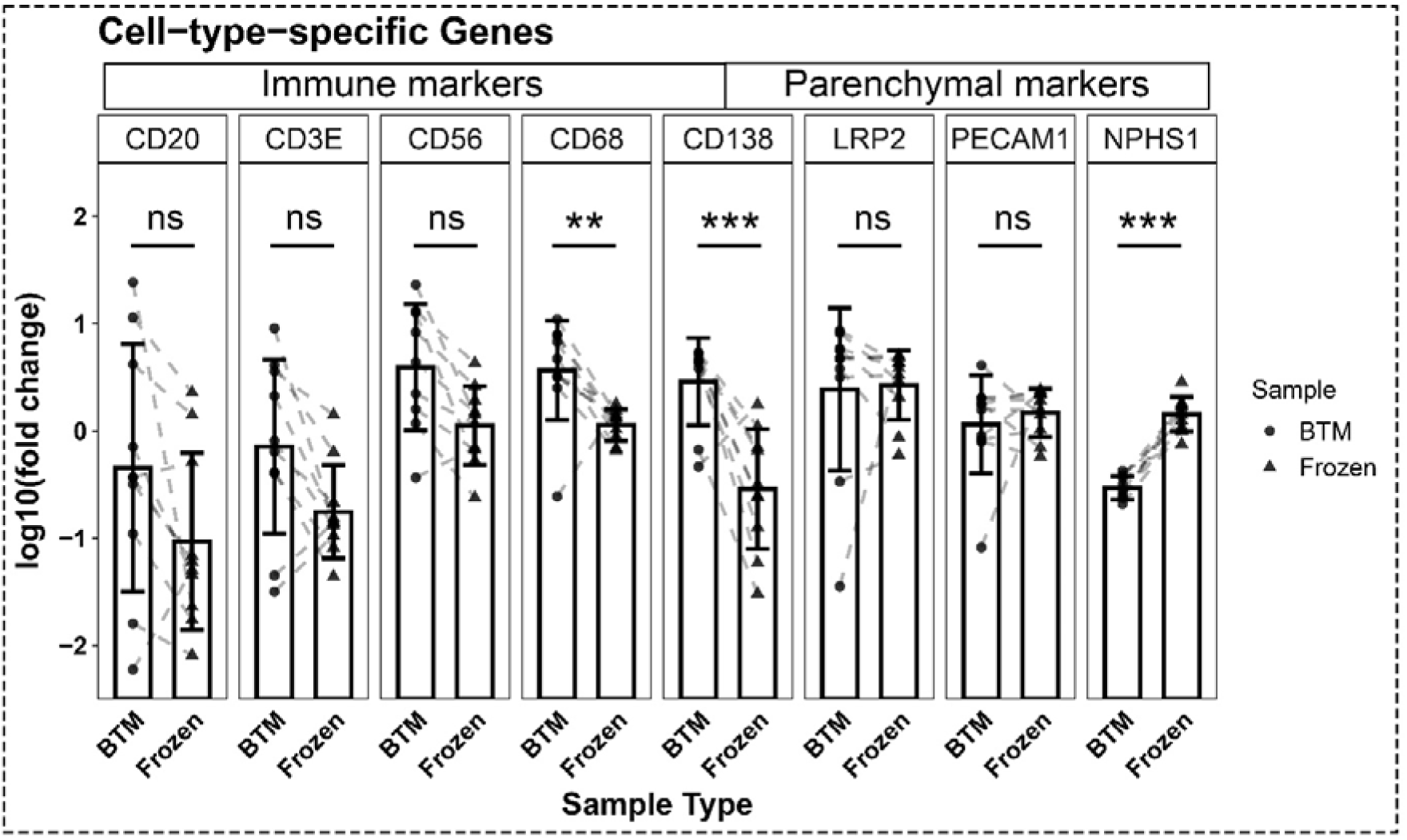
Cell distribution in mimicked BTM relative to paired adjacent frozen tissue. Expression of immune and parenchymal cell markers was quantified in both BTM samples and their corresponding frozen tissues. Each dot in the Frozen group denotes an individual kidney, whereas each dot in the BTM group represents the mean expression value across all BTM samples derived from the same kidney. Lines connected the samples derived from the same kidneys. The y-axis depicts expression levels normalized to *HPRT1*.

### 3.5 Reliability and agreement analysis between mimicked BTM and the corresponding fresh biopsies, frozen tissues, and FFPE tissues

For RNA-Seq, 9 samples were created, consisting of 3 mimicked biopsy tissues, 3 BTM, and 3 adjacent frozen tissues derived from 3 nephrectomies. For NanoString B-HOT panel analysis, 12 samples were created, comprising 7 BTM (of these, 5 derived from the same kidney but at different storage time points, and the other two from two separate kidneys), 3 adjacent FFPE tissues, and 2 adjacent frozen tissues derived from 3 nephrectomies (Table 2). In total of 6 comparison groups were established based on sample types: 3 groups based on RNA-Seq data (BTM vs Fresh biopsy tissues, BTM vs Frozen tissues, Fresh biopsy tissues vs Frozen tissues) and 3 groups based on NanoString data (BTM vs FFPE tissues, BTM vs Frozen tissues, and FFPE tissues vs Frozen tissues). The quantification of reliability analysis yielded ICC values for the above 6 groups, which were, respectively, 0.84, 0.84, 0.90, 0.82, 0.88, and 0.90. High reliability was observed in all comparison groups, with ICC values exceeding 0.80 (Table 3). Bland-Altman plots analysis was performed on log10-transformed data to assess agreement in gene expression between 6 comparison groups. Mean bias ranged from –100.41 to 100.21, corresponding to a 0.39- to 1.62-fold difference between methods. The 95% limits of agreement varied across groups, ranging from 0.03- to 9.33-fold. Significant proportional bias was observed in all groups, with slopes equivalent to 0.72- to 1.66-fold change (all p < 0.001). (Table 4, Figure 4). Although agreement was partly dependent on expression magnitude, most biases were close to 1-fold, suggesting that overall agreement between sample types was good.

**Figure 4.**
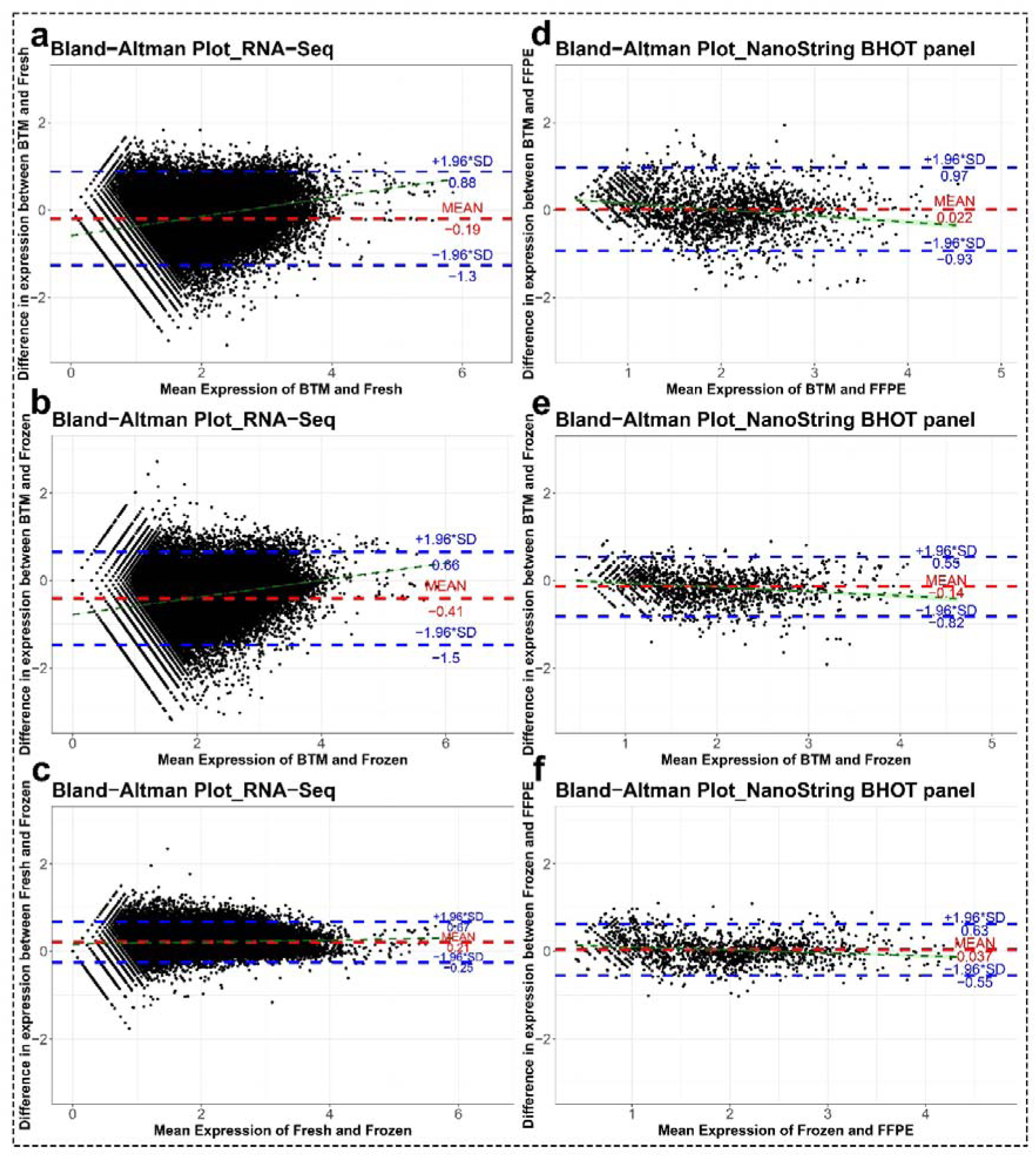
Bland-Altman plots illustrating the agreement in gene expression between the samples from the same kidney but different processing types. (a,b, and c) comparisons based on RNA-Seq data, and (d, e, and f) comparisons based on NanoString B-HOT panel. The y-axis represents the difference between methods, while the x-axis represents their mean. The red lines denote the mean difference, indicating any systematic bias, and the blue lines indicate the 95% limits of agreement (mean difference ± 1.96 SD). The fitted regression line (dark green) (with 95% confidence interval shading (light green)) shows whether the difference between methods varies systematically across the measurement range. The analysis is based on a log10 scale; most data points lie within these limits, suggesting that the two methods show acceptable agreement.

**Table 2.**
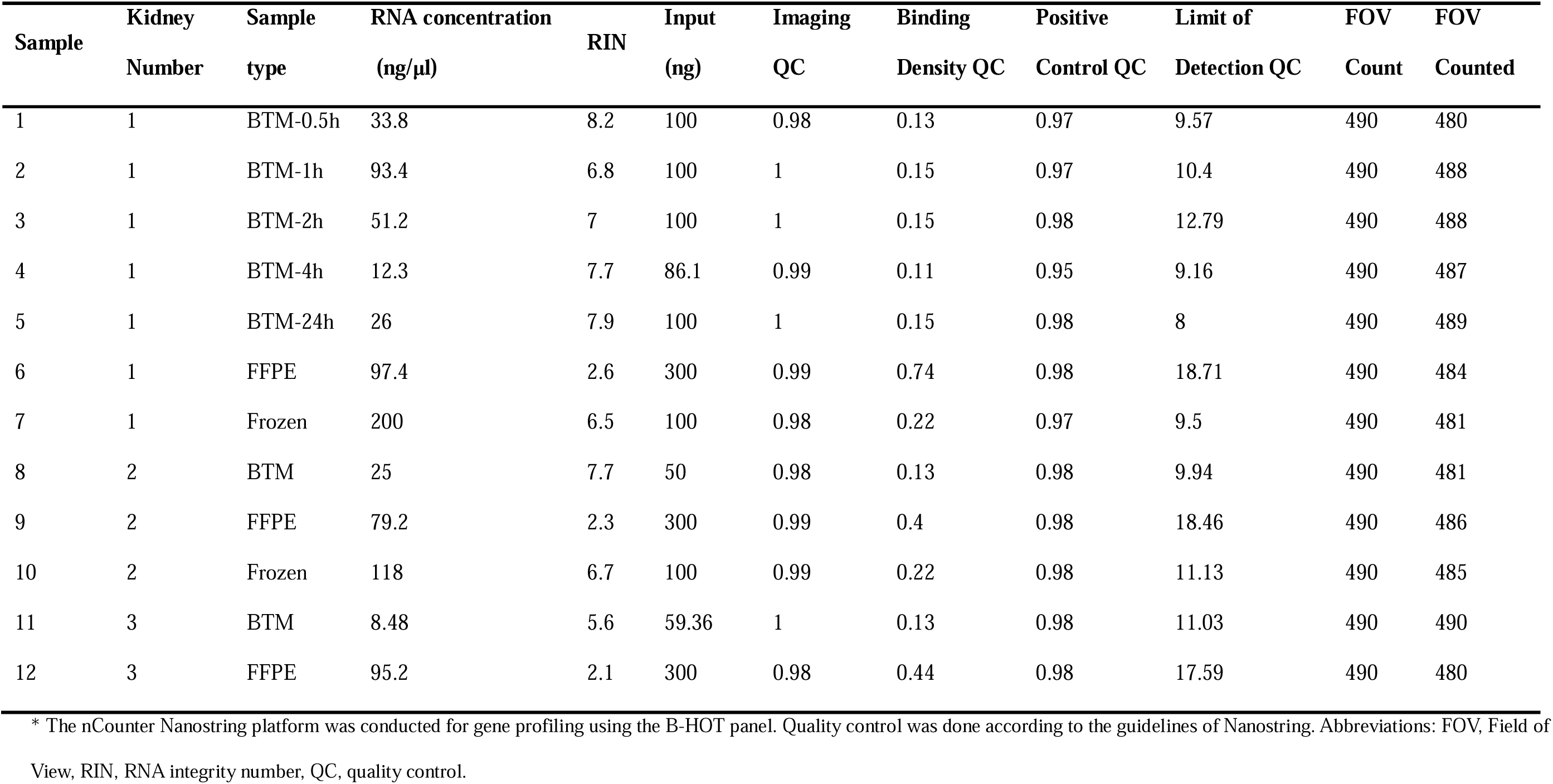
Characteristics of the samples selected for the comparison of different time points of BTM versus the corresponding FFPE and frozen material.

| Sample | Kidney Number | Sample type | RNA concentration (ng/μl) | RIN | Input (ng) | Imaging QC | Binding Density QC | Positive Control QC | Limit of Detection QC | FOV Count | FOV Counted |
| --- | --- | --- | --- | --- | --- | --- | --- | --- | --- | --- | --- |
| 1 | 1 | BTM-0.5h | 33.8 | 8.2 | 100 | 0.98 | 0.13 | 0.97 | 9.57 | 490 | 480 |
| 2 | 1 | BTM-1h | 93.4 | 6.8 | 100 | 1 | 0.15 | 0.97 | 10.4 | 490 | 488 |
| 3 | 1 | BTM-2h | 51.2 | 7 | 100 | 1 | 0.15 | 0.98 | 12.79 | 490 | 488 |
| 4 | 1 | BTM-4h | 12.3 | 7.7 | 86.1 | 0.99 | 0.11 | 0.95 | 9.16 | 490 | 487 |
| 5 | 1 | BTM-24h | 26 | 7.9 | 100 | 1 | 0.15 | 0.98 | 8 | 490 | 489 |
| 6 | 1 | FFPE | 97.4 | 2.6 | 300 | 0.99 | 0.74 | 0.98 | 18.71 | 490 | 484 |
| 7 | 1 | Frozen | 200 | 6.5 | 100 | 0.98 | 0.22 | 0.97 | 9.5 | 490 | 481 |
| 8 | 2 | BTM | 25 | 7.7 | 50 | 0.98 | 0.13 | 0.98 | 9.94 | 490 | 481 |
| 9 | 2 | FFPE | 79.2 | 2.3 | 300 | 0.99 | 0.4 | 0.98 | 18.46 | 490 | 486 |
| 10 | 2 | Frozen | 118 | 6.7 | 100 | 0.99 | 0.22 | 0.98 | 11.13 | 490 | 485 |
| 11 | 3 | BTM | 8.48 | 5.6 | 59.36 | 1 | 0.13 | 0.98 | 11.03 | 490 | 490 |
| 12 | 3 | FFPE | 95.2 | 2.1 | 300 | 0.98 | 0.44 | 0.98 | 17.59 | 490 | 480 |

\* The nCounter Nanostring platform was conducted for gene profiling using the B-HOT panel. Quality control was done according to the guidelines of Nanostring. Abbreviations: FOV, Field of View, RIN, RNA integrity number, QC, quality control.

**Table 3.**
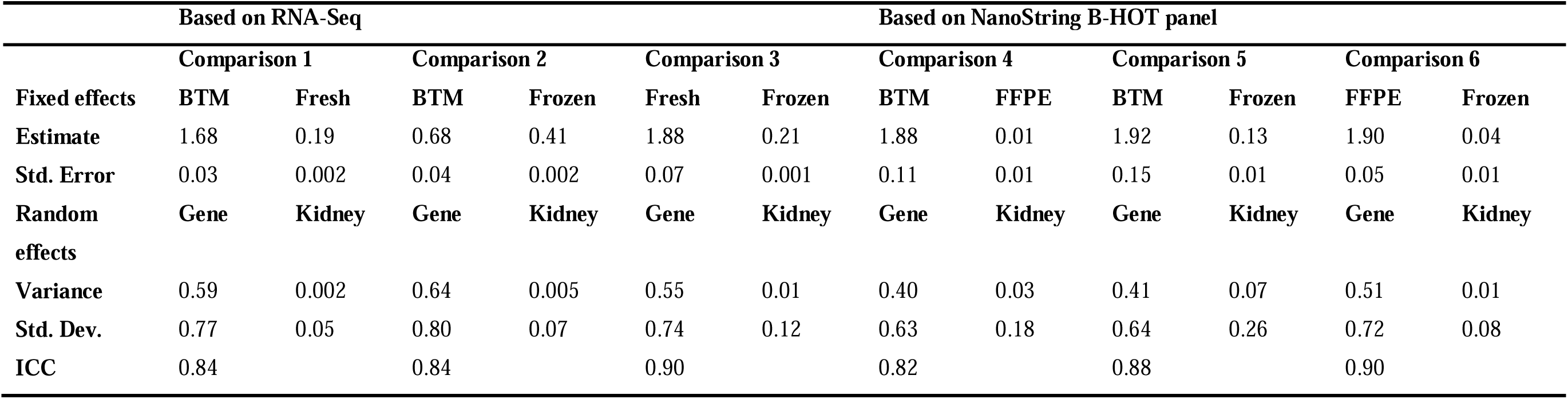
Interclass Correlation Coefficient (ICC) Analysis (Std. Error = Standard Error, Std. Dev. = Standard Deviation)

**Table 4.** Summary of Bland–Altman Analysis Across Six Comparison Groups (LoA = limits of agreement)

|  | Comparison | Mean Difference (log10) | 95% LoA (log10) | Slope (Proportional Bias, log10) | p-value |
| --- | --- | --- | --- | --- | --- |
| Based on RNA-Seq | BTM vs Fresh | -0.19 | -1.27, 0.88 | 0.22 | <0.001 |
|  | BTM vs Frozen | -0.41 | -1.47, 0.66 | 0.20 | <0.001 |
|  | Fresh vs Frozen | 0.21 | -0.25, 0.67 | 0.02 | <0.001 |
| Based on NanoString | BTM vs FFPE | 0.02 | -0.93, 0.97 | -0.14 | <0.001 |
| B-HOT panel | BTM vs Frozen | -0.14 | -0.82, 0.55 | -0.10 | <0.001 |
|  | FFPE vs Frozen | 0.04 | -0.55, 0.63 | -0.07 | <0.001 |

### 3.6 Cell composition estimation on mimicked BTM and the corresponding fresh biopsies and frozen tissues

Cell fractions were estimated by deconvoluting bulk RNA-seq data using CIBERSORTx. Across all samples, the proportions of immune and parenchymal cells showed variability among different sample types (Figure 5a). At the level of individual cell populations, all cell populations detected in fresh biopsy tissues were also identified in the corresponding BTM samples based on deconvolution cell fraction estimates (Figure 5b).

**Figure 5.**
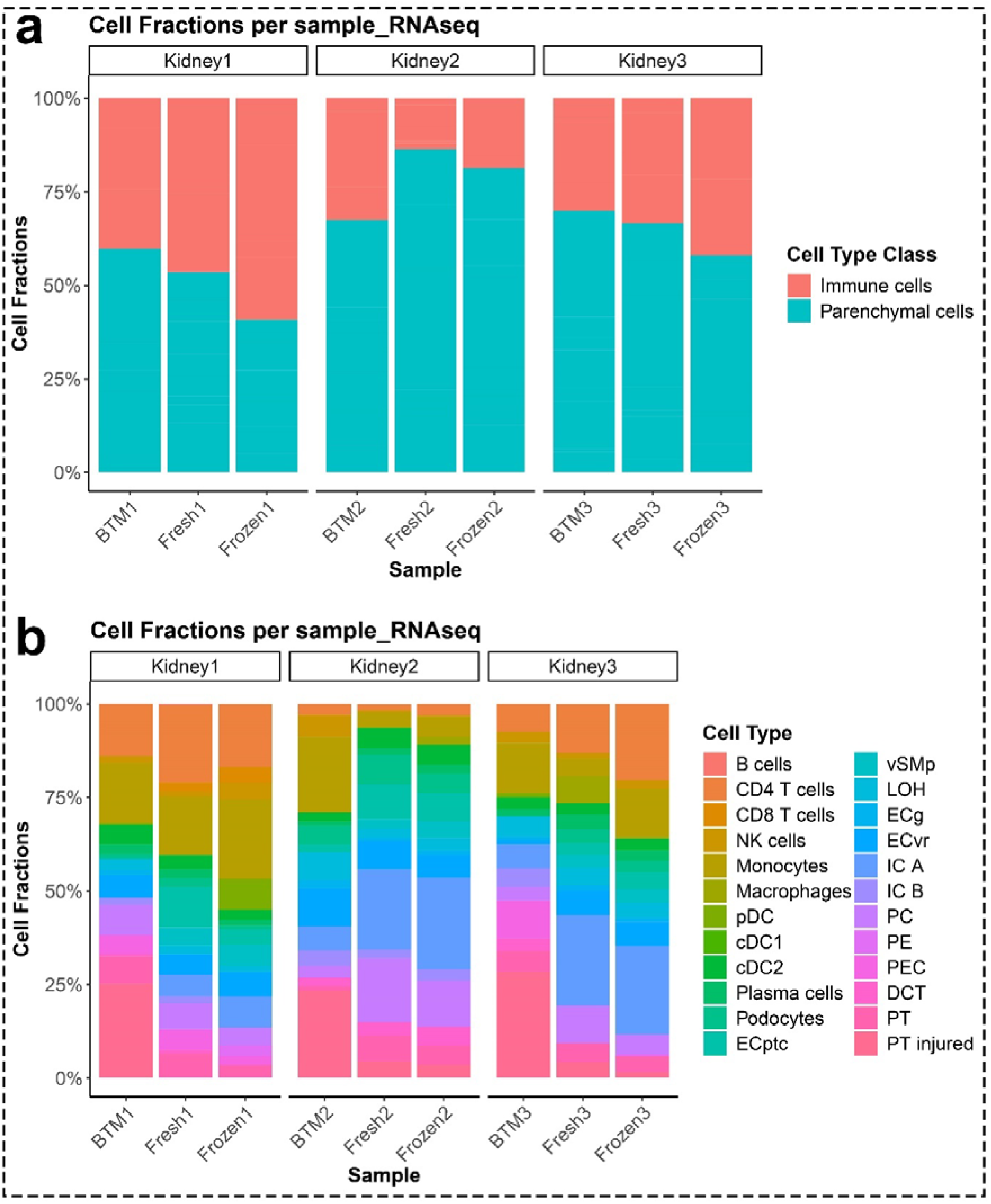
Cell type composition estimated from RNA-Seq data using cellular deconvolution. (a) Stacked bar plot shows the relative proportions of immune cells and parenchymal cells across all samples, with each bar representing an individual sample and the y-axis indicating proportional abundance. (b) The same samples are shown with each major cell class further resolved into its concrete types. Each bar represents one sample, with stacked segments corresponding to the estimated contribution of each cell type.

DGE analysis was performed on NanoString B-HOT panel data. Between BTM and the corresponding FFPE tissues, a total of 98 genes were higher in BTM, while 92 genes were higher in FFPE (adjusted p-value < 0.05, |log2FC| > 1) (Figure 6a). To refine the analysis, we focused on genes with more pronounced differences (adjusted p-value < 0.05, |log2FC| > 2.5), identifying 68 differentially expressed genes (DEGs). Mapping these genes to kidney single-cell type clusters (Human Protein Atlas^7^) revealed that the BTM group displayed a higher proportion of immune cell gene expression compared to FFPE (74% vs. 15%), while the FFPE group exhibited a higher proportion of genes expressed in parenchymal cells compared to BTM (27% vs. 9%) (Figure 6b). These results corroborate previous qPCR results and cell fractions results, indicating enrichment of immune cell transcripts in the BTM compared to the corresponding biopsy tissue.

**Figure 6.**
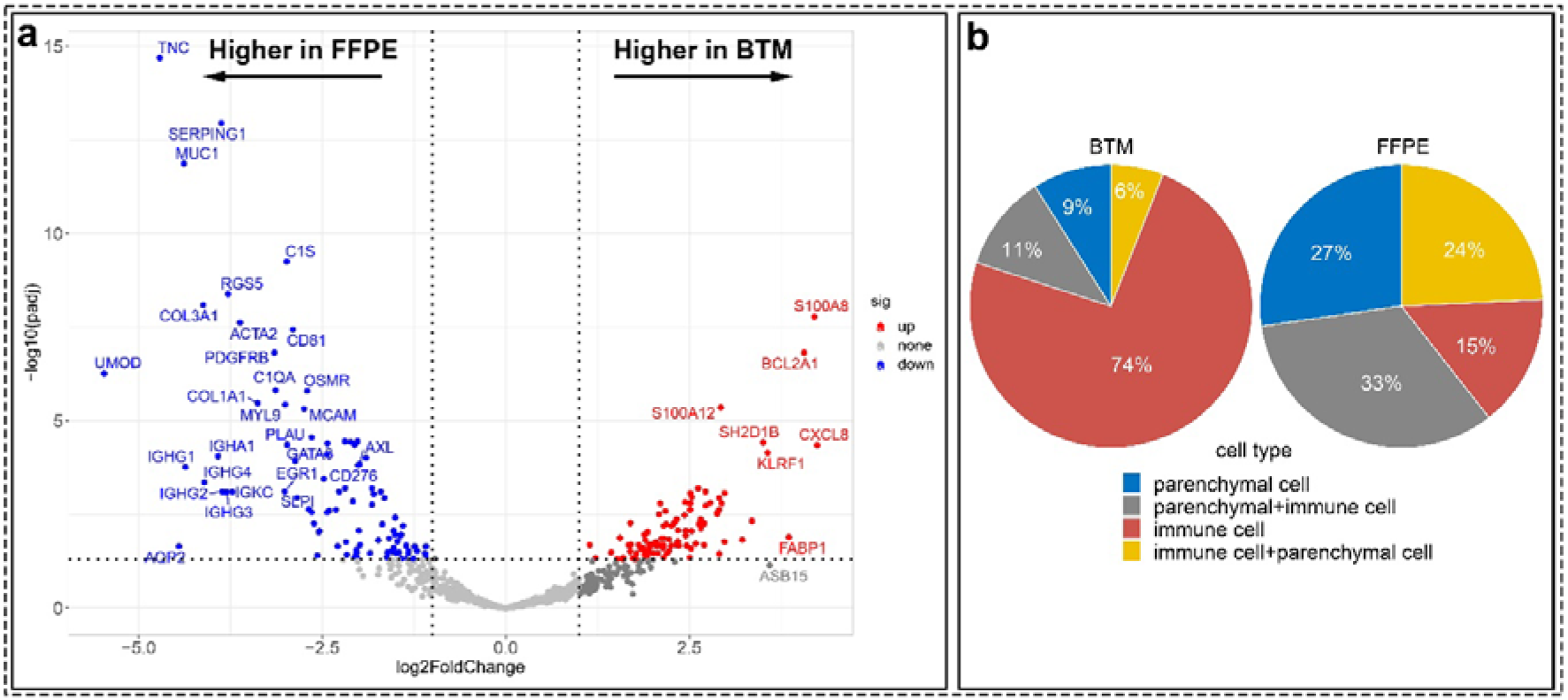
The difference in gene distribution between the mimicked BTM and the corresponding FFPE. (**a**) The volcano plot depicts B-HOT panel gene expression in BTM (kidney 1 (mean of five storage time points), kidney 2, and kidney 3) and FFPE (kidney 1, kidney 2, and kidney 3). Each colored dot represents a gene. The fold change (FC) is the ratio of mRNA count in BTM to FFPE. The X-axis represents the log2FC values. The 2 dotted vertical lines on either side of 0 on the X-axis represent |log2FC| = 1. The Y-axis represents the -log10 adjusted P value. The horizontal dotted line on the Y-axis represents adjusted p-value=0.05. The blue and red dots are genes that are |log2FC| > 1 and have an adjusted p-value < 0.05. Thus, red dots represent statistically significant DEGs that are higher in BTM compared with FFPE, and the blue dots represent the relatively higher levels in FFPE. Of the 770 genes, 98 were higher and 92 were lower in BTM. (**b**) Pie charts show the proportion of different cell types that are enriched by 68 overexpressed DEGs(|log2FC| > 2.5). Parenchymal cell + immune cell means mRNA expressed in both cell types, but parenchymal cells account for the majority. Immune cell + parenchymal cell means mRNA expressed in both cell types, but immune cells account for the majority.

### 3.7 Proof-of-concept pilot study on the real transplant BTM to determine clinical potential

Kidney transplant biopsies taken in the context of DGF represent a real-world clinical scenario in which fast molecular profiling could reduce the time to tailored treatment. In DGF, the clinical differential diagnosis is often between acute tubular necrosis (ATN), related to the deceased donation procedure, and requiring supportive treatment only, or rejection, which would benefit from timely anti-rejection treatment. Twelve random anonymized BTM samples from clinical kidney transplant biopsies with DGF were used for NanoString B-HOT panel measurement to investigate whether gene expression profiling can discriminate between these clinical scenarios (Table 5). Given the exploratory nature of this pilot study, we prioritized identifying DEGs with a less stringent criterion of P-value < 0.05, which ensures that potential gene candidates were not overlooked. The analysis showed a total of 54 genes were differentially expressed (p-value < 0.05 and |log2FC| > 1) between histologically proven ATN and rejection (Antibody-Mediated Rejection (AMR) and T Cell-Mediated Rejection (TCMR) together), with 23 genes exhibiting higher expression in the ATN group and 31 genes exhibiting higher expression in the rejection group (Figure 7a). The heatmap of the DGEs further provides preliminary evidence that samples within the rejection group clustered into AMR and TCMR (Figure 7b). Further DGE analysis between AMR and TCMR identified 49 genes (p-value < 0.05 and |log2FC| > 1) (Figure 7c), with 12 genes higher in the TCMR group and 37 genes higher in the AMR group (Figure 7d).

**Figure 7.**
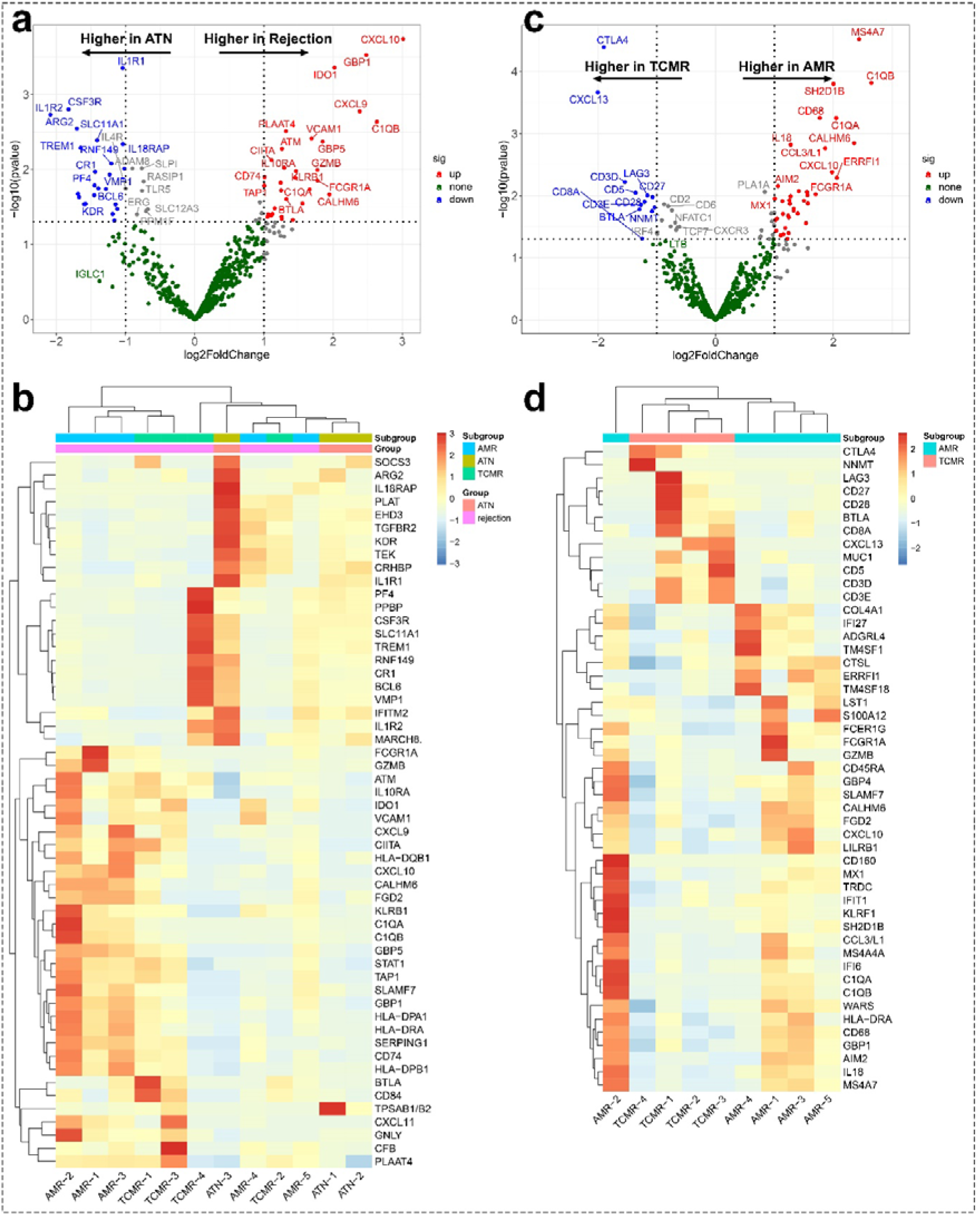
Differentially expressed genes (DEGs) on 12 real BTM. Two volcano plots (a and c) depict B-HOT panel gene expression, with each dot representing a gene. The fold change (FC) is the ratio of mRNA count between two groups. The X-axis represents the log2FC values. The 2 dotted vertical lines on either side of 0 on the X-axis represent |log2FC| = 1. The Y-axis represents the -log10 P value. The horizontal dotted line on the Y-axis represents P-value = 0.05. The blue and red dots are genes that have |log2FC| > 1, respectively, and a P-value < 0.05. Two heatmaps (b and d) show the top DEGs, with each column representing a BTM sample and each row representing a gene. (a) The red dots represent statistically significant genes that are higher in rejection compared to ATN, and the blue dots represent the genes that are higher in ATN. (b) Hierarchical clustering suggests significant heterogeneity between rejection and ATN. Of the 770 genes, 31 were higher and 23 were lower in rejection. (c) The red dots represent statistically significant genes that are higher in AMR compared to TCMR, and the blue dots represent the genes that are higher in TCMR. (d) Hierarchical clustering suggests significant heterogeneity between AMR and TCMR. Of the 770 genes, 37 were higher and 12 were lower in AMR.

**Table 5.** Characteristics of the samples selected for the proof of principle run of clinical samples on Nanostring and the results of the Nanostring Quality Control (RIN= RNA integrity number)

| Sample | Diagnosis | RNA concentration<br>(ng/μl) | RIN | Input (ng) | Imaging<br>QC | Binding<br>Density QC | Positive<br>Control QC | Limit of<br>Detection QC | FOV<br>Count | FOV<br>Counted |
| --- | --- | --- | --- | --- | --- | --- | --- | --- | --- | --- |
| 1 | AMR-1 | 5.54 | 6.3 | 38.78 | 0.99 | 0.17 | 0.99 | 6.12 | 490 | 485 |
| 2 | AMR-2 | 4.4 | 3.1 | 30.8 | 1 | 0.2 | 0.99 | 9.12 | 490 | 490 |
| 3 | AMR-3 | 5.64 | 4.8 | 39.48 | 1 | 0.19 | 0.98 | 7.62 | 490 | 480 |
| 4 | AMR-4 | 7.88 | N/A | 55.16 | 0.99 | 0.24 | 0.98 | 8.38 | 490 | 480 |
| 5 | AMR-5 | 8.62 | 8.6 | 60.34 | 0.99 | 0.25 | 0.96 | 6.75 | 490 | 486 |
| 6 | TCMR-1 | 7.9 | 6.8 | 55.3 | 1 | 0.19 | 0.98 | 7.12 | 490 | 488 |
| 7 | TCMR-2 | 6 | 5.9 | 42 | 1 | 0.16 | 0.98 | 7.5 | 490 | 488 |
| 8 | TCMR-3 | 10.4 | 5.9 | 72.8 | 0.99 | 0.2 | 0.99 | 9.88 | 490 | 487 |
| 9 | TCMR-4 | N/A | 6.1 | N/A | 0.99 | 0.14 | 0.97 | 6.64 | 490 | 489 |
| 10 | ATN-1 | 8.64 | 8.1 | 60.48 | 0.98 | 0.21 | 0.98 | 7 | 490 | 484 |
| 11 | ATN-2 | 7.08 | 4.8 | 49.56 | 1 | 0.18 | 0.98 | 8.19 | 490 | 481 |
| 12 | ATN-3 | 5.48 | 4 | 38.36 | 1 | 0.14 | 0.98 | 6.12 | 490 | 481 |

\* The nCounter Nanostring platform was conducted for gene profiling using the B-HOT panel. Quality control was done according to the guidelines of Nanostring. Abbreviations: AMR, antibody-mediated rejection; TCMR, T cell-mediated rejection; ATN, acute tubular necrosis; FOV, Field of View, RIN; N/A, not available; RNA integrity number; QC, quality control.

We further investigated the correlation between gene expression in BTM samples and the histologic findings based on the corresponding FFPE biopsies. The frequency of each Banff lesion score across the corresponding FFPE biopsies is summarized in Supplementary Figure 8. Per-lesion Spearman correlation analyses were performed, correlating the expression of differentially expressed genes identified in the AMR-versus-TCMR comparison with individual Banff lesion scores (Figure 8). This analysis revealed distinct associations between transcript expression level and specific histopathologic lesions, with several genes showing significant positive correlations, highlighting coordinated molecular changes linked to lesion severity. The genes higher in the AMR group, such as *CD160, CD68, CXCL10, GBP1, MS4A7*, and *SH2D1B*, demonstrated positive correlations with lesion score g (glomerulitis), which is a feature of activity and antibody interaction with tissue in AMR (ρ > 0.7, p < 0.05). In contrast, the genes higher in TCMR group, such as *BTLA, CD3D*, and *CD3E*, were positively correlated with lesion score i (interstitial inflammation), t (tubulitis), and ti (total inflammation), which are key diagnostic features of TCMR in kidney transplants (ρ > 0.6, p < 0.05). Additionally, *CD8A*, higher in the TCMR group, showed a moderate correlation with ti (ρ > 0.6, p < 0.05), and *CTLA4*, higher in the TCMR group, was positively correlated with both i and t (ρ > 0.6, p < 0.05).

**Figure 8.**
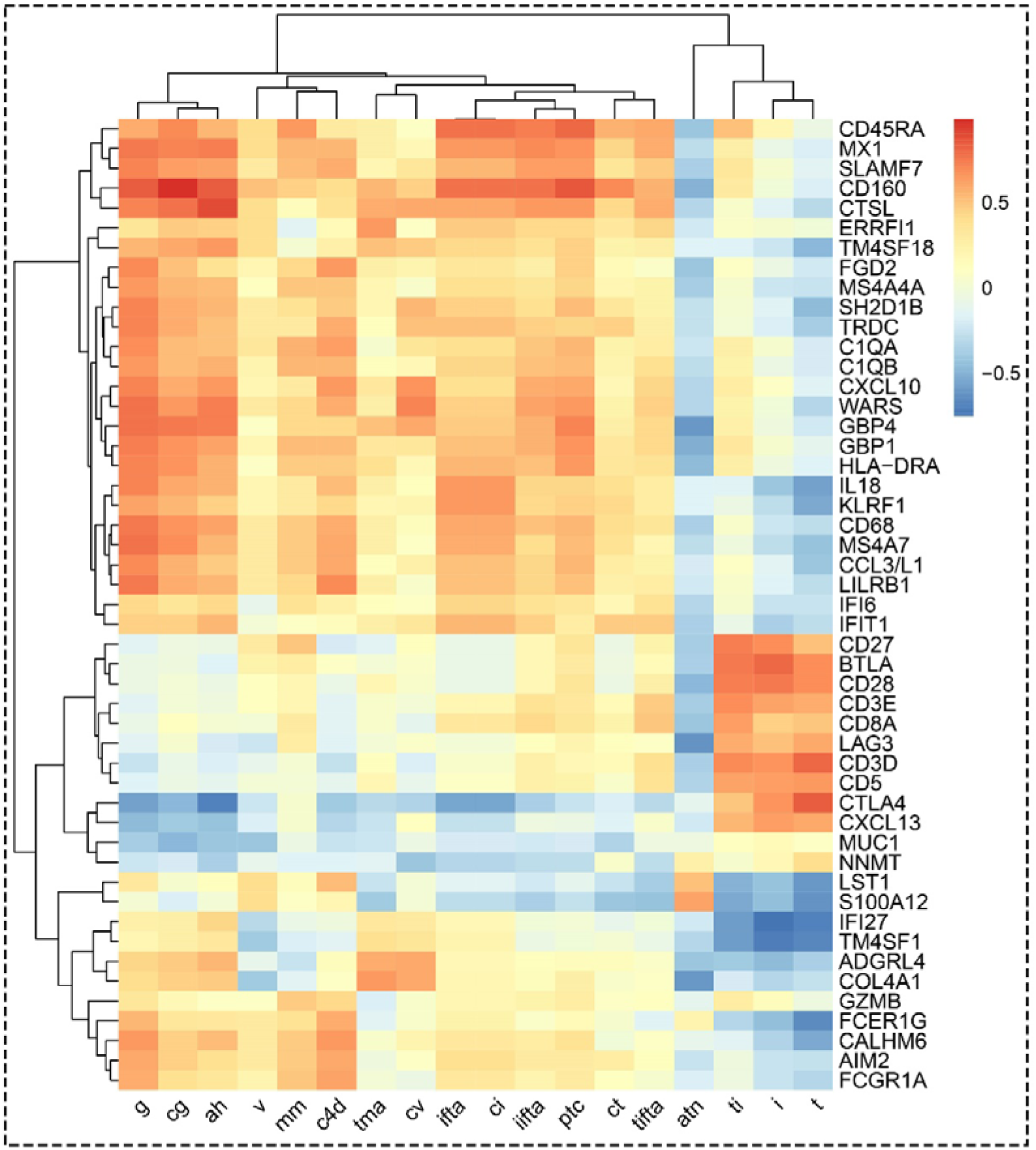
Heatmap of Spearman’s rank correlation coefficients comparing the DEGs with Banff lesion scores in real BTM in the setting of DGF. Each cell represents the Spearman correlation coefficient between paired observations (DEGs and Banff lesion scores) across samples. The analysis aims to assess whether changes observed at the gene level follow the same trend as those seen with Banff lesion scores. Red colors indicate positive correlations, while blue colors indicate negative correlations; deeper colors represent stronger correlation coefficients. White or lighter shades reflect weak or no correlation. Correlation coefficients range from –1 (perfect negative) to +1 (perfect positive). atn, acute tubular necrosis; ti, total interstitial inflammation; i, interstitial inflammation; t, tubulitis; tifta, tubulitis in the area of IFTA; iifta, inflammation in the area of IFTA; ptc, peritubular capillaritis; tma, thrombotic microangiopathy; ct, tubular atrophy; ifta, interstitial fibrosis and tubular atrophy; ci, interstitial fibrosis; v, intimal arteritis; g, glomerulitis; cg, GBM double contours; ah, arteriolar hyalinosis; mm, mesangial matrix expansion; cv, vascular fibrous intimal thickening.

## 4 DISCUSSION

Our study establishes BTM as a novel and yet unexplored liquid biopsy with potential utility for molecular diagnosis. We optimized and validated the technical methods for RNA isolation and assessed the effect of storage time, determining that samples processed within 24 hours qualified for multiple transcriptomic technologies. Additionally, in a proof-of-concept study, we demonstrated the potential for BTM gene expression profiling to reflect the status of transplanted kidneys, consistent with the Banff lesion score. These findings need further exploration in prospective observational studies.

Since BTM originates from routine biopsy procedures yet has rarely been examined as a biological material, our first objective was to confirm the presence of viable biological information within this medium. Our preliminary experiments confirmed the presence of cells in the BTM using flow cytometry and cytospin, with cytospin-prepared morphology consistent with biopsy findings. This consistency across methodologies reinforces the robustness of our observations and provides a critical foundation for subsequent RNA-based analyses. The mechanisms by which cells are released into the BTM merit careful consideration. We hypothesize two principal pathways: first, direct mechanical detachment during the biopsy procedure, in which shear stress or tissue disruption liberates cells into the surrounding medium; and second, cell release during the post-collection preservation phase, potentially due to enzymatic activity or loss of cellular adhesion. Importantly, our data revealed no significant differences in RNA yield across different storage time points. This finding suggests that mechanical forces during biopsy may perhaps be the predominant factor contributing to cellular release, while post-preservation processes may play a minor role.

Despite the inherent instability of RNA, our study demonstrated that RNA extracted from BTM remains of sufficient integrity for downstream molecular applications provided that samples are processed within 24 hours under appropriate storage conditions. This is critical observation, as it indicates that BTM is compatible with transcriptomic analyses, thereby extending the potential use of biopsy-derived materials beyond conventional tissue samples aiding fast diagnosis. In terms of RNA isolation protocol, the TRIzol-based method was limited by small starting materials, which frequently resulted in minimal or invisible RNA precipitation. This makes it challenging to get rid of residual phenol or guanidine, consequently increasing the risk of contamination. In contrast, the RNeasy Kit provided more reliable and user-friendly workflow, yielding RNA of higher purity and consistency and making it the preferred method for RNA isolation from BTM.

The reliability and agreement analyses demonstrate that BTM is not only a technically reliable source for gene expression analysis but also a biologically meaningful one, capable of reflecting the transcriptional landscape of the tissue of origin. Both RNA-Seq and NanoString B-HOT panel results, BTM showed strong concordance with corresponding fresh biopsy tissues or FFPE tissues across a broad spectrum of genes. Cellular deconvolution further revealed that BTM recapitulates the cell populations present in fresh biopsy tissues, with only subtle differences. All major cell types present in the tissue were detectable in BTM, although there was a slightly higher proportion of immune cells and a marginally lower proportion of parenchymal cells. These findings are expected since parenchymal cells are anchored to each other and to basement membranes and/or collagen fibers through tight and adherent junctions, while immune cells are more prone to migration out of the needle biopsy. The targeted B-HOT panel analysis on 12 real BTM derived from kidney transplant biopsies further reinforces the translation relevance. The rejection-associated genes identified in BTM closely matched biomarkers previously reported in FFPE and conventional biopsy specimens^8^, indicating that BTM can recapitulate clinically relevant molecular signatures. Moreover, the observed correlation of AMR- and TCMR-associated gene expression with their respective Banff lesion scores provides additional validation that molecular alterations captured in BTM reflect underlying histopathological changes.

As a novel liquid-based biological resource, our findings highlight a key advantage of BTM over urine in its ability to capture endothelial cell–derived signals that are absent from urine due to the anatomical localization of endothelial cells. This observation is corroborated by a recent single-cell RNA sequencing study of urine, in which endothelial cells were not detected^9^. In our analyses, the detection of *PECAM1* and other endothelial-enriched genes such as *PLAT* supports the potential of BTM for monitoring endothelial injury and its potential relevance in AMR, where such injury is central^10^. Podocyte-associated transcripts such as *NPHS1* were markedly lower in BTM than in frozen tissues; however, as the samples were not from patients with podocytopathies^11^. This likely reflects cohort characteristics rather than limitations of BTM, underscoring the need for future studies in cohorts with podocyte injury.

The ultimate goal of our research is to enable a rapid molecular assessment in settings with a limited diagnostic window. While not universally applicable, we confirmed that transporting biopsies in PBS is sufficient to preserve tissue integrity without compromising subsequent analyses. In this regard, BTM emerges as an attractive supplementary liquid biopsy resource that could precede or complement conventional histological evaluation once the biopsy reaches the pathology department. Such an approach could provide particular value in urgent clinical scenarios, such as renal allografts at risk of acute rejection, which significantly affects allograft survival^12,13^. By facilitating earlier recognition of molecular changes, BTM may support the initiation of biologically targeted interventions to mitigate graft injury at an earlier stage and thereby improve long-term outcomes^14^. An additional strength of BTM lies in its accessibility and practicality. Because it represents a by-product of the biopsy transport process, its collection requires no additional sampling or tissue processing, distinguishing it from most other molecular diagnostic platforms. This “low-burden” nature of BTM collection enhances its feasibility for routine clinical integration. Furthermore, by providing rapid and objective molecular readouts, BTM has the potential to overcome some of the limitations of conventional microscopy-based assessments, which rely heavily on expert interpretation and are subject to inter-observer variability.

In conclusion, our results demonstrate that RNA obtained from BTM is of sufficient quality for gene expression analysis. BTM represents a promising new liquid biopsy material for rapid molecular assessment in cases with narrow diagnostic windows. This approach is not restricted to kidney biopsies and may be optimized for any fresh biopsies transported to pathology. Future research will focus on developing targeted gene panels for primary discrimination between clinically relevant differential diagnoses to guide initial treatment choices and prevent further organ damage in case of aggressive disease presentations.

## Abbreviation

AKI: Acute Kidney Injury
MMDx: Molecular Microscope Diagnostic System
B-HOT: Banff Human Organ Transplant
FFPE: Formalin-Fixation, Paraffin-Embedding
PBS: Phosphate-Buffered Saline
BTM: Biopsy Transport Medium
LUMC: Leiden University Medical Center
DGF: Delayed Graft Function
SOP: Standard Operating Procedure
PI: Propidium Iodide
RIN: RNA Integrity
Ct: Quantification Cycle
HPRT1: Hypoxanthine phosphoribosyltransferase 1
FACS: Fluorescence-Activated Cell Sorting
RNA-Seq: RNA Sequencing
QC: Quality Control
DGE: Differential Gene Expression
ICV: Inter-assay Coefficient of Variance
SD: Standard Deviations
ICC: Intraclass Correlation Coefficient
TPM: Transcript-Per-Million
UBB: Ubiquitin B
OAZ1: Ornithine decarboxylase antizyme 1
PPIA: Peptidylprolyl isomerase A
ATN: Acute Tubular Necrosis
DEGs: Differentially Expressed Genes
AMR: Antibody-Mediated Rejection
TCMR: T Cell-Mediated Rejection

## Acknowledgment

The study is supported by the Department of Pathology, Leiden University Medical Center and the China Scholarship Council (CSC). We would like to acknowledge Ron Wolterbeek (Department of Biomedical Data Sciences, Medical Statistics, Leiden University Medical Centre, Leiden, The Netherlands) for support.

## Data statement

We have deposited the RNA-Seq data and NanoString data sets generated for this study at the Gene Expression Omnibus (https://www.ncbi.nlm.nih.gov/gds/) with accession numbers GSE308751, GSE274322, and GSE274323.

## Supplementary document

### S1 MATERIALS and METHODS

#### S1.1 Preliminary experiments

##### S1.1.1 Cytospin Preparation and morphology assessment on real Biopsy Transport Medium (BTM)

Resuspend the white pellet in 500 µl PBS buffer. Loaded the cell suspension into cytospin chambers and centrifuged at 1,500 rpm for 5 minutes using Cytospin 4 cytocentrifuge (Thermo Scientific, Waltham, MA, USA). After air-drying, the standard microscope slide was stained with Giemsa stock solution (Sigma Aldrich 1.09204.0500, Germany) following standard protocols.

##### S1.1.2 Flow cytometry on real BTM

Resuspend the white pellet and wash the cells twice with Fluorescence-Activated Cell Sorting (FACS) buffer (PBS containing 0.5% bovine serum albumin). Two markers were selected: *CD45*, a pan-leukocyte marker expressed on nearly all hematopoietic cells, and *LRP2* (megalin), a multiligand endocytic receptor abundantly expressed on the apical surface of proximal tubular epithelial cells. Then, the cells were stained with BV421-labeled Anti-Human *CD45* antibody (1:200, BioLegend; 368521) and an unconjugated primary anti-*LRP2* antibody (1:500, Leiden University Medical Center, Leiden, the Netherlands) followed by Alexa Fluor® 647 goat anti-mouse IgG antibody (1:400, Thermo Fisher; A-21244). Cells were incubated sequentially with primary and secondary antibodies at 4 °C for 30 minutes with gentle shaking, and washed three times with FACS buffer between each incubation step. Live/dead cell discrimination was performed using propidium iodide (PI) just before acquisition on a BD FACS Canto-II 3L flow cytometer (BD Biosciences, San Jose, CA, USA).

### S1.2 Effect of ultracentrifugation speed on mimicked BTM

To examine whether an ultracentrifugation speed will affect the gene expression analysis, three extra mimicked BTM were created and equally divided into two parts: one centrifuged at 1,500g and the other at 20,000g. The RNA isolation procedure remained identical. The difference in housekeeping gene expression was analyzed.

### S1.3 RNA quantity and integrity assay

RNA concentration was measured with the NanoDrop 2000 spectrophotometer (Thermo Scientific, Wilmington, DE, USA) and the Qubit RNA HS Assay Kit on the Qubit 2.0 Fluorometer (Thermo Fisher Scientific, Waltham, MA, USA) according to the manufacturer’s instructions. RNA integrity (RIN) was assayed using the Agilent 2100 Bioanalyzer System(Agilent Technologies, Santa Clara, CA, USA).

### S1.4 Hematoxylin and eosin (H&E) staining on tumor-free kidney FFPE tissue sections

Formalin-fixed paraffin-embedded (FFPE) tissue sections (derived from the corresponding kidneys, which were used to create mimicked BTM for procedural comparison) were cut at a thickness of 4 μm and mounted on glass slides. Then, the slides were deparaffinized, rehydrated, and stained with H&E following standard protocols.

### S1.5 Bulk RNA-sequencing (RNA-Seq) analysis in mimicked BTM and the corresponding biopsies and frozen tissues

The mimicked BTM, the corresponding fresh kidney biopsy, and the adjacent frozen tissue were used for bulk RNA-sequencing. RNA-Seq FASTQ files were processed using the open-source BIOWDL RNAseq pipeline v5.0.0 developed in LUMC. This pipeline performs FASTQ preprocessing (including quality control, quality trimming, and adapter clipping), RNA-Seq alignment and read expression quantification. FastQC was used for checking raw read QC. Adapter clipping was performed using Cutadapt (v2.10) with default settings. RNA-Seq reads alignment was performed using STAR (v2.7.5a) on the GRCh38 human reference genome. The gene read quantification was performed using HTSeq-count (v0.12.4) with the setting ‘–stranded=yes’. The gene annotation used for quantification was Ensembl version 104. Genes with counts greater than 10 in at least two samples were included in downstream analysis.

### S1.6 NanoString B-HOT panel expression analysis in mimicked and real BTM

Since FFPE tissue is the most widely used specimen type in conventional diagnostic practice but is suboptimal for RNA-seq due to extensive RNA degradation, we performed gene expression profiling using the NanoString nCounter MAX Analysis System (NanoString Technologies, Seattle, WA, USA). The hybridization-based NanoString platform is inherently compatible with FFPE material, enabling reliable gene expression profiling and spatial analysis from archival or fragmented samples. The B-HOT code set, comprising 770 genes, was used to analyze gene expression of RNA isolated from mimicked BTM, the corresponding frozen and FFPE tissues, as well as real BTM. Hybridization of RNA samples with the probes occurred at 65°C for 17 hours, followed by loading onto the nCounter Prep Station in accordance with the manufacturer’s instructions. Following data collection with the nCounter MAX Digital Analyzer, a preliminary analysis was conducted using the nSolver 4.0 Analysis Software System, including quality control (QC) based on Imaging QC, Binding Density QC, Positive Control QC, and Limit of Detection QC. Samples that passed QC were subjected to a two-step data transformation process (positive control normalization and housekeeping gene normalization), and normalized counts were exported for subsequent differential gene expression (DGE) analysis.

### S1.7 Statistical analysis

IBM SPSS Statistics 29.0.0.0 software was used for the comparative and agreement analysis. The differences in RNA yield across five storage time groups (RNA yield [ng] = RNA concentration [ng/μl] × volume [μl]) were examined through one-way ANOVA. The RIN values of RNA isolated by different procedures were compared with the Independent-Samples T-test. To explore the effects of storage times, centrifuge speed, and RNA isolation methods on RNA quantity, *HPRT1* Ct values were analyzed using one-way ANOVA or Independent-Samples T-test. Column bar graphs and connected dot plots were generated by GraphPad Prism 9.3.1(471). The agreement of gene expression between BTM versus the corresponding fresh biopsies, FFPE, and frozen tissues was assessed by computing intraclass correlation coefficient (ICC), which was calculated from linear mixed effects statistical models with random effects of kidneys and genes. The appropriateness of the models was examined visually by histograms of the residuals from each of the two models. The corresponding visualization was displayed via Bland-Altman plots. The DGE analysis and the corresponding visualization were implemented by RStudio-4.2.3 in Windows 11. DGE analysis utilized the “DESeq2-1.38.3” package (downloaded from https://bioconductor.org/packages/release/bioc/html)”. The heatmaps were generated with the ‘pheatmap-1.0.12’ package”. The volcano plots were generated with the ‘tidyverse-2.0.0’ and ‘ggrepel-0.9.4’ packages. The Bland-Altman Plots were generated with the ggplot2-3.4.4’ packages. CIBERSORTx was used to deconvolute the predicted cell fractions from the bulk RNA-Seq data. The ‘biomaRt-2.62.1’ package was used to preprocess gene count data into transcript-per-million (TPM) space. The signature matrix used in this study was derived from single-cell RNA sequencing of 16 kidney allograft biopsies and enables the discrimination of 24 distinct parenchymal and immune cell phenotypes

### S2 FIGURES

**Supplementary Figure 1.**
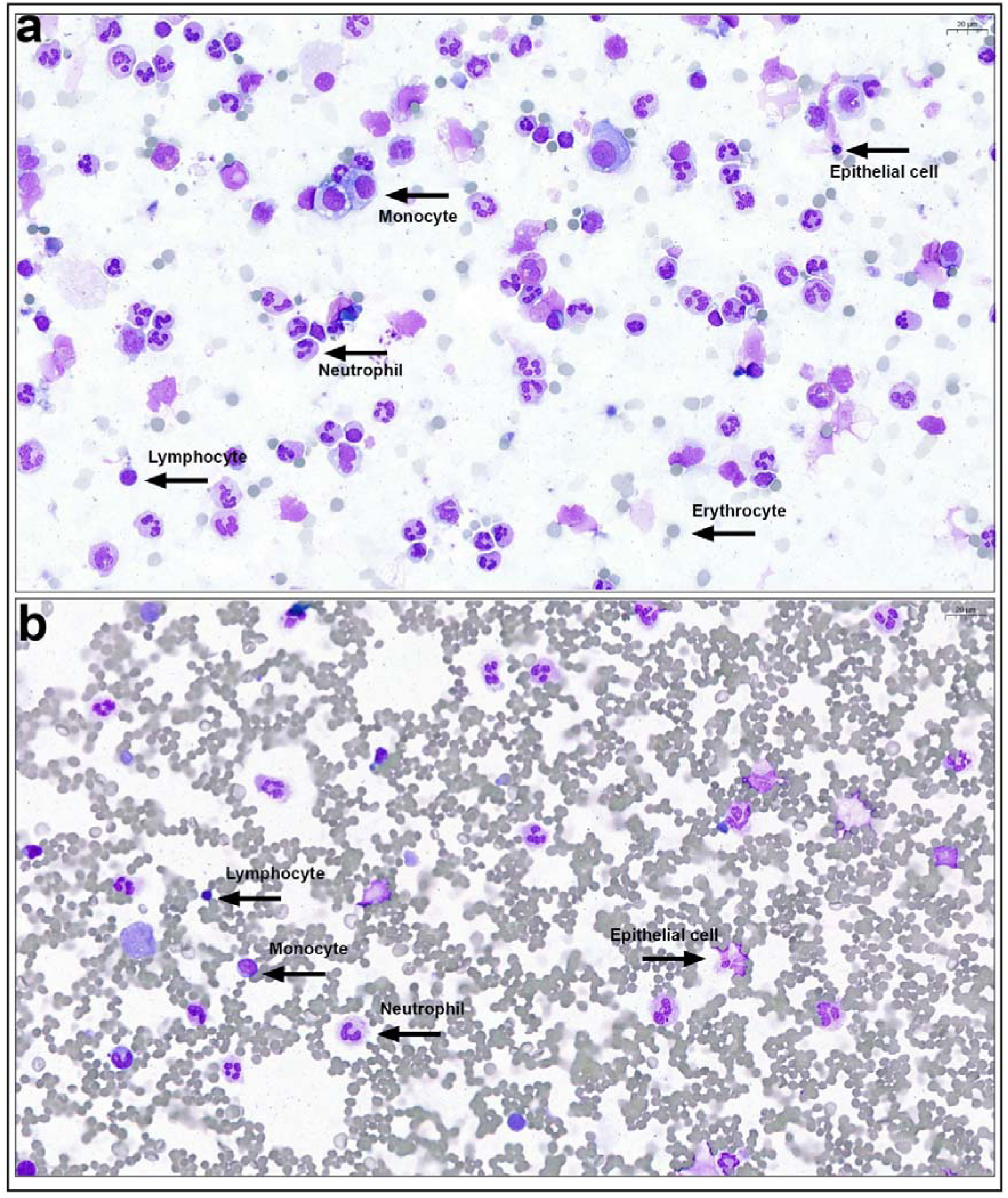
Wright–Giemsa staining of cytospin-prepared samples. The other two BTM samples demonstrate well-preserved cellular morphology, enabling clear identification of distinct cell populations.

**Supplementary Figure 2.**
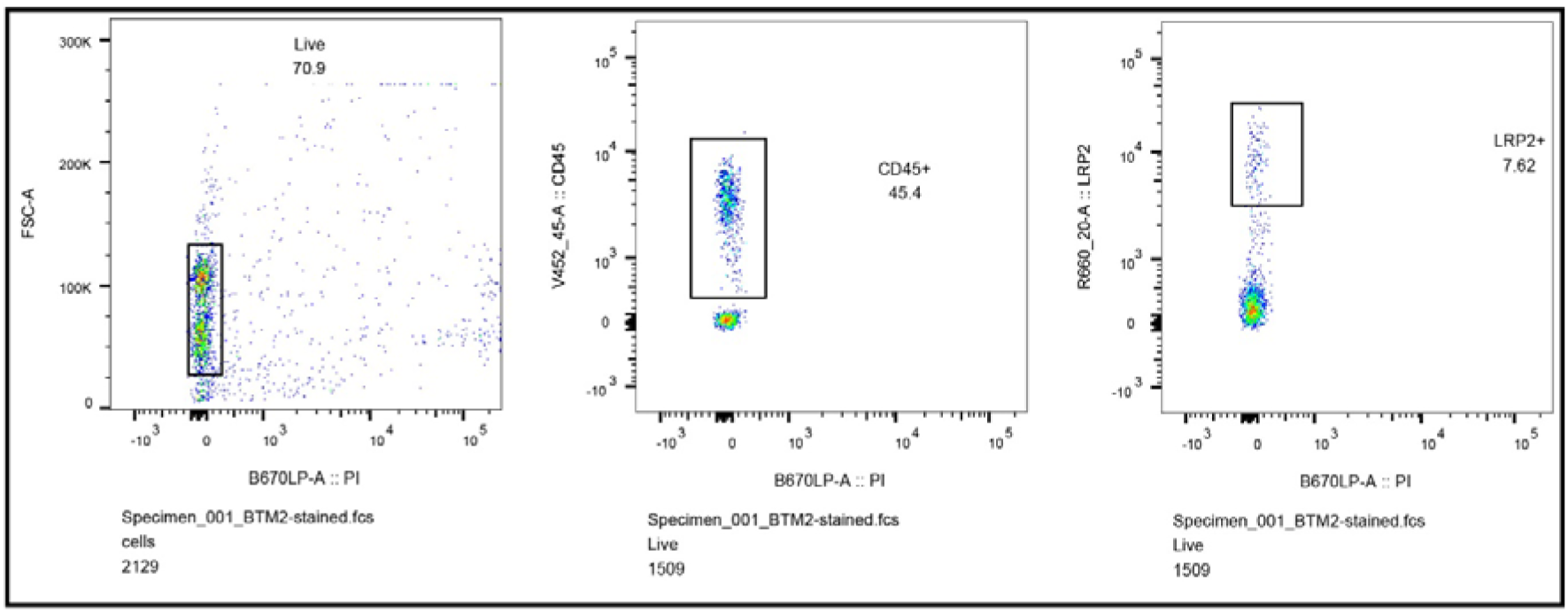
Flow cytometry analysis of CD45 and LRP2 expression. Flow cytometry analysis was performed to evaluate the expression of CD45, indicative of immune cells, and LRP2, representing epithelial lineage, on a real BTM sample. After gating on living cells (71%), and excluding debris, the expression profiles of CD45 and LRP2 were assessed. CD45L cells constituted 45% of the total live cell population, while LRP2L cells accounted for 7.6% of the total live cell population.

**Supplementary Figure 3.**
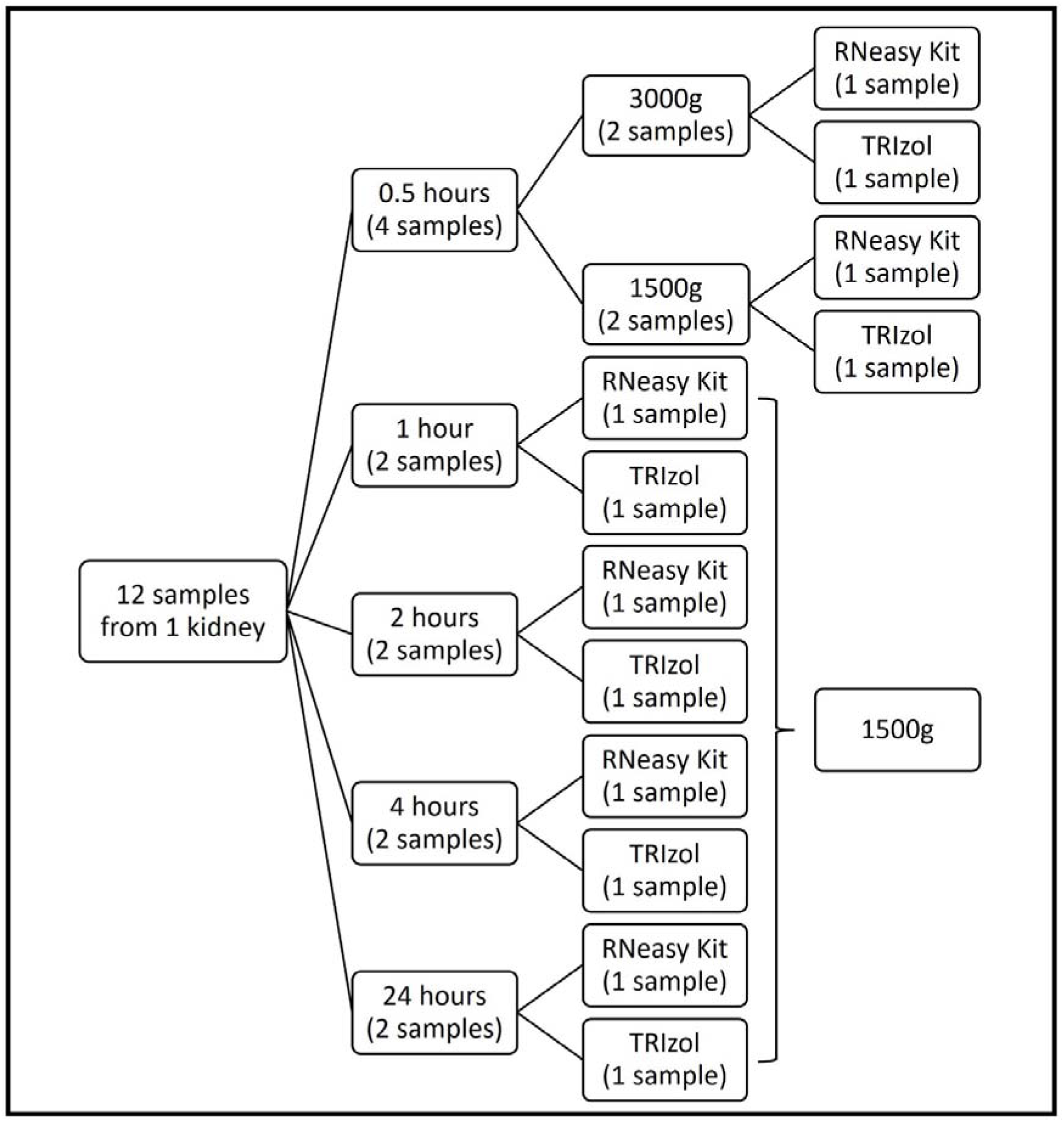
Workflow of procedural comparisons. Three factors that potentially affect the RNA yield were tested: storage times, centrifuge speed, and RNA isolation methods. Each sample consists of 2 biopsies with 15 ml of PBS. There are 12 samples in total, 2 samples were assigned to each storage time point group, except for the 0.5 hours group, to which 4 samples were assigned. Then, the two samples of each storage time point group, except for the 0.5 hours group, were divided into 2 groups (one that was isolated by the TRIzol method and the other one by RNeasy Kit) after centrifuging at the speed of 1500g. The four samples of the 0.5 hours group were divided into 2 groups, one was centrifuged at the speed of 3000g, and another at the speed of 1500g. Following that, 2 samples in each speed group were equally divided into 2 RNA isolation method groups.

**Supplementary Figure 4.**
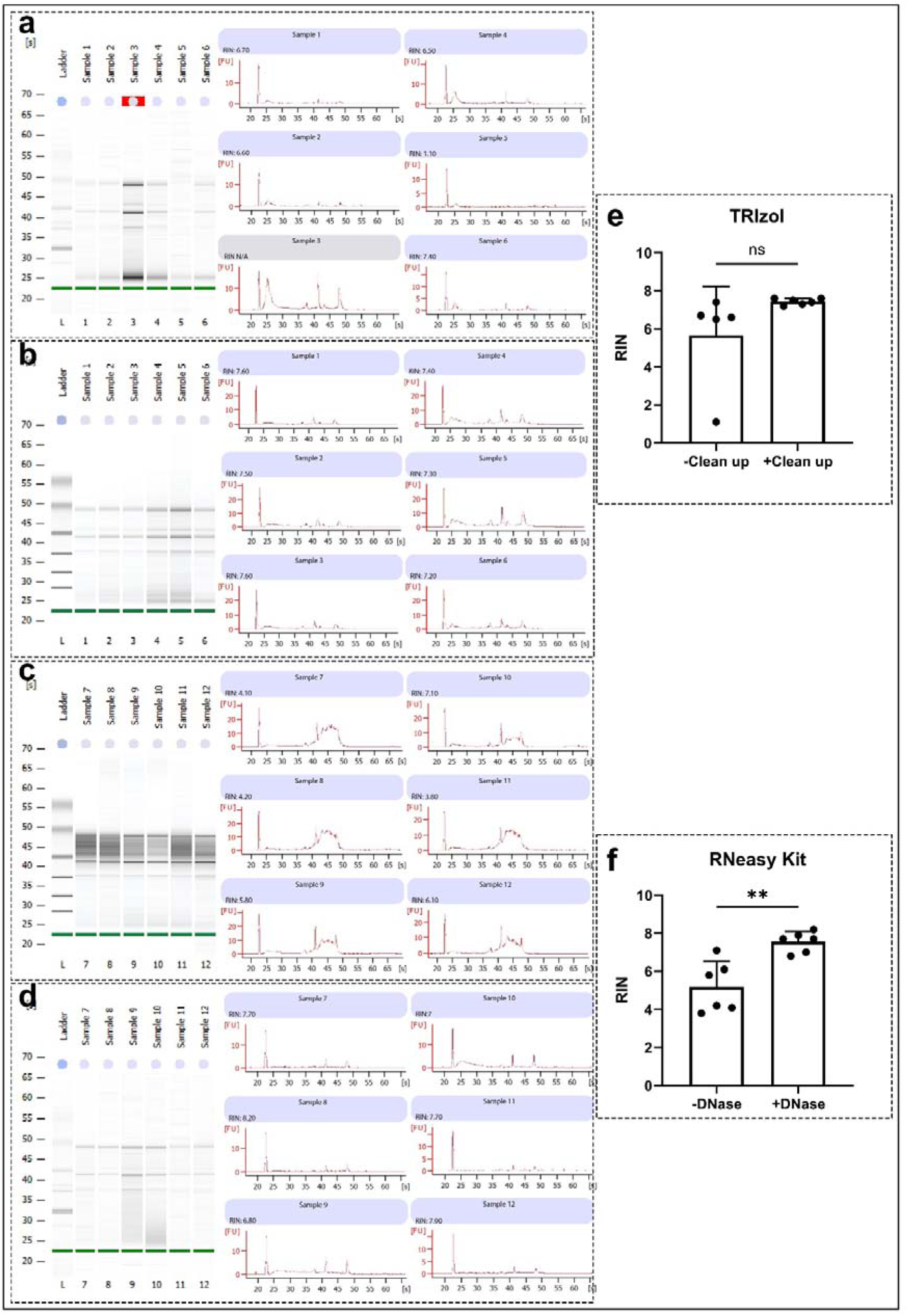
The RNA integrity assay. Two distinct and sharp bands in the gel image correspond to the 28S (upper band) and 18S (lower band) rRNA, indicating intact RNA. The electropherogram displays fluorescence intensity (y-axis) versus migration time or fragment size (x-axis), and the two main peaks represent the 18S and 28S ribosomal RNA. (a) The RNA was isolated from 6 samples using only the TRIzol method. Sample 3 shows an “N/A” warning, indicating that the RIN may be unreliable. Sample 5 does not show a high-intensity band in the gel image and has an extremely low RIN value of 1.1. (b) The RNA was isolated from another 6 samples, which were derived from the same kidney, using the TRIzol method plus the RNeasy clean up. This method shows higher RIN values with less variation after the RNeasy clean-up procedure compared to TRIzol alone. (c) The RNA was isolated from 6 samples using RNeasy without DNase treatment. The presence of dark backgrounds and broad peaks between 18S and 28S suggests contamination with genomic DNA. (d) RNA was isolated from another 6 samples, derived from the same kidney, using RNeasy with DNase treatment. This method shows clear bands in the gel image and higher RIN values after DNase treatment compared to the previous results. Two bar charts represent mean RIN ± standard deviation for each group. Individual dots represent individual samples. (e) Comparison of RIN between TRIzol alone (-clean up) and TRIzol method plus the RNeasy clean up (+clean up); (f) Comparison of RIN between RNeasy without DNase treatment (-DNase) and RNeasy with DNase treatment (+DNase).

**Supplementary Figure 5.**
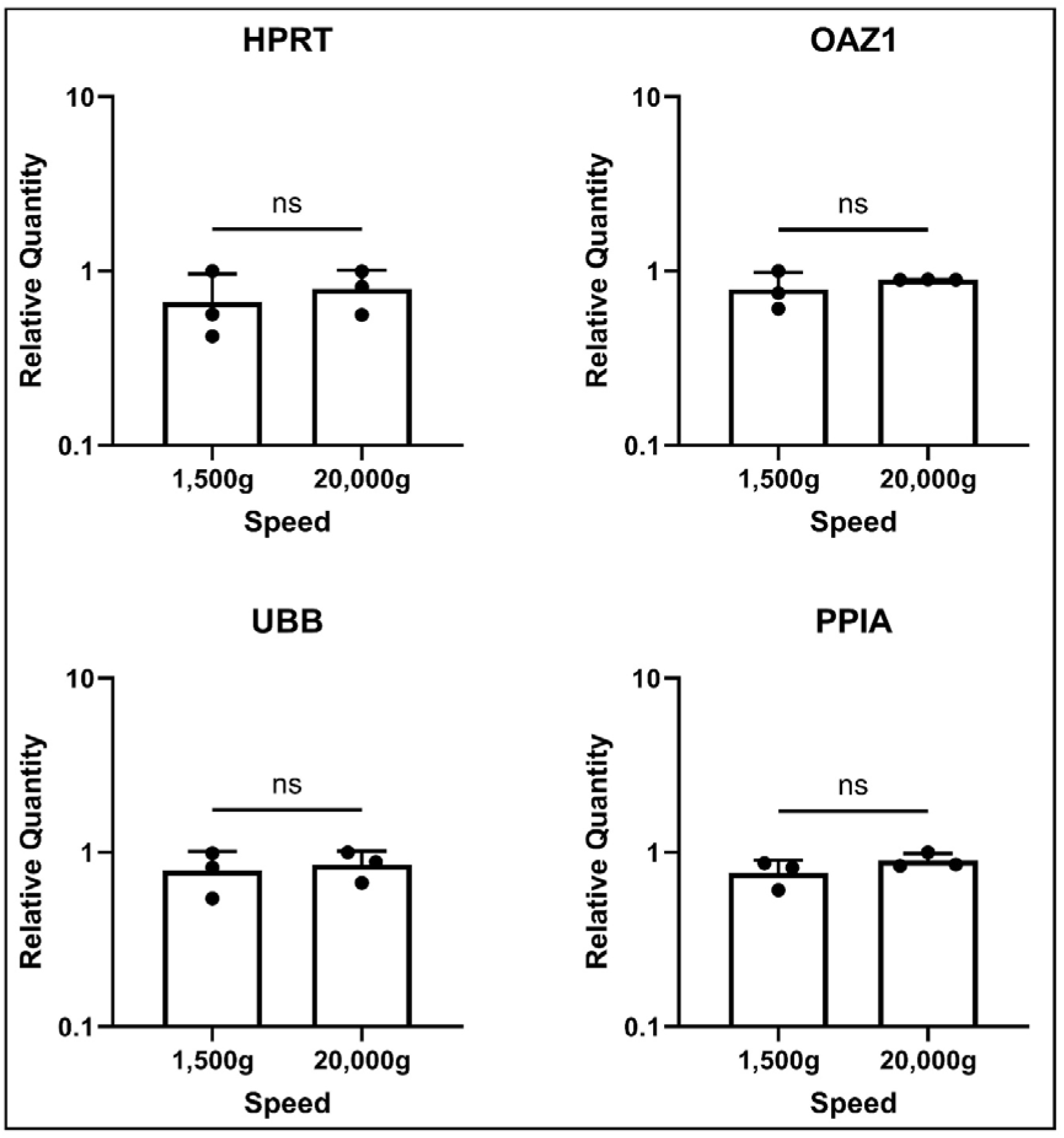
Relative quantities of four housekeeping genes across comparison groups. Bars represent the mean relative quantity of genes within each group; individual dots indicate the relative quantity of individual samples. Error bars represent SD. Groups correspond to different centrifugation speeds. Three mimicked BTM samples were involved, and the expression of four housekeeping genes (HPRT1, Ubiquitin B (UBB), Ornithine decarboxylase antizyme 1 (OAZ1), and Peptidylprolyl isomerase A (PPIA)) was measured.

**Supplementary Figure 6.**
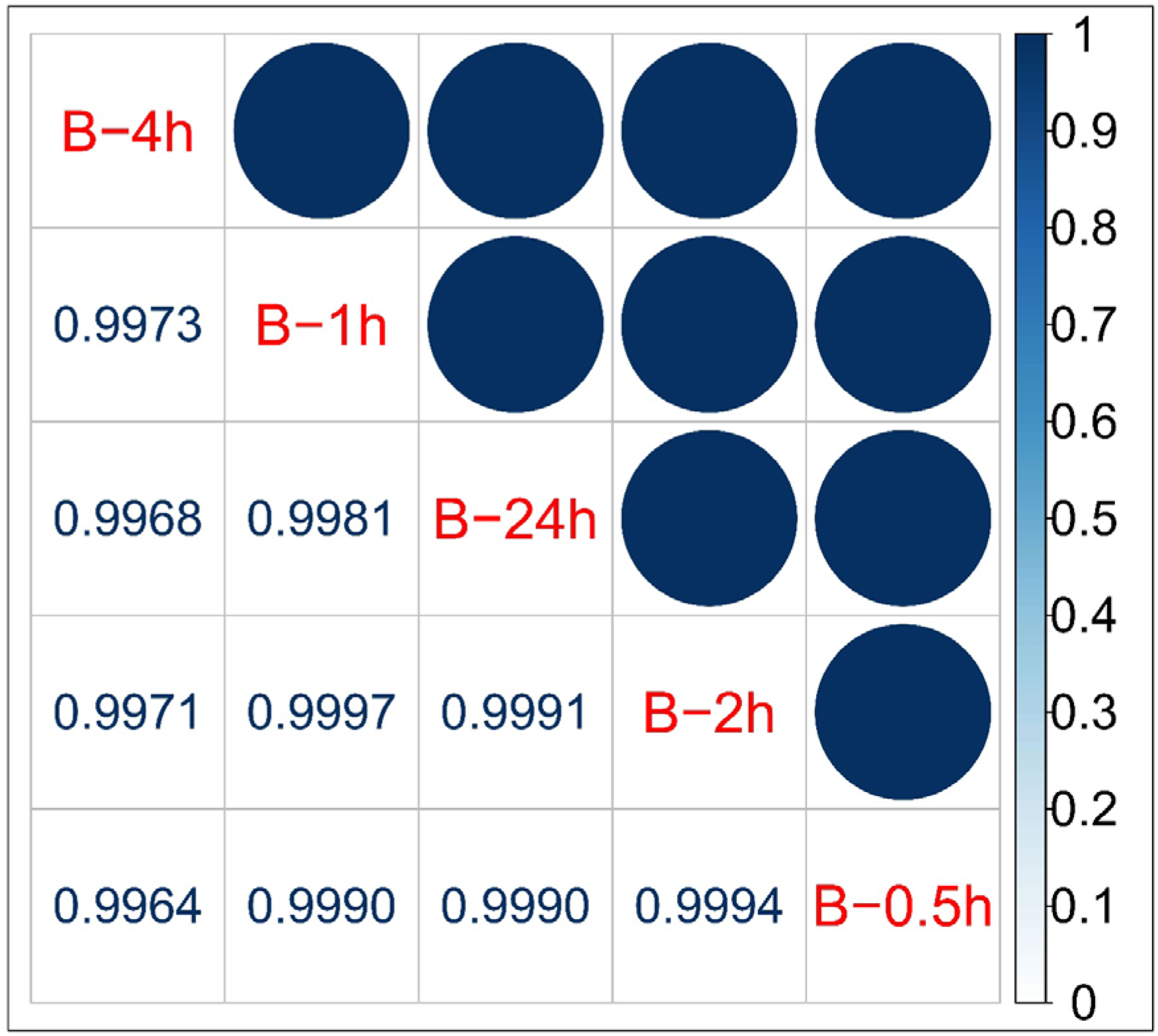
Correlation matrix for NanoString B-HOT expression among mimicked BTM storage times. A strong correlation between the gene expression of the B-HOT panel among the five storage time points (all R >0.99).

**Supplementary Figure 7.**
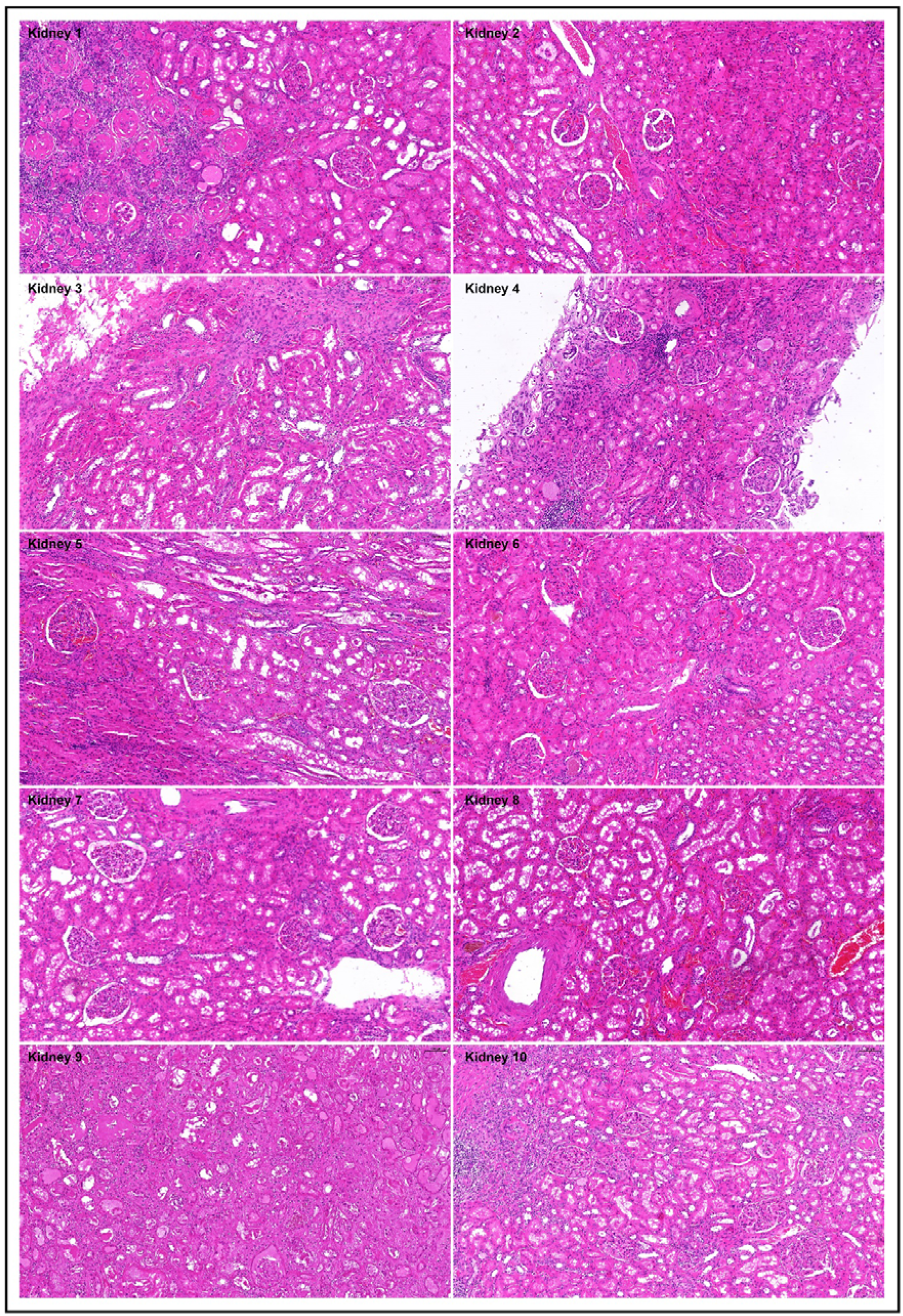
H&E staining of FFPE kidney tissue. Adjacent tissues from the same 10 nephrectomies were processed into FFPE tissues for histological analysis. Digital slide evaluation revealed focal areas of inflammation, characterized by immune cell infiltration within the tissues. Scale bar: 200 µm, at the upper right of each figure.

**Supplementary Figure 8.**
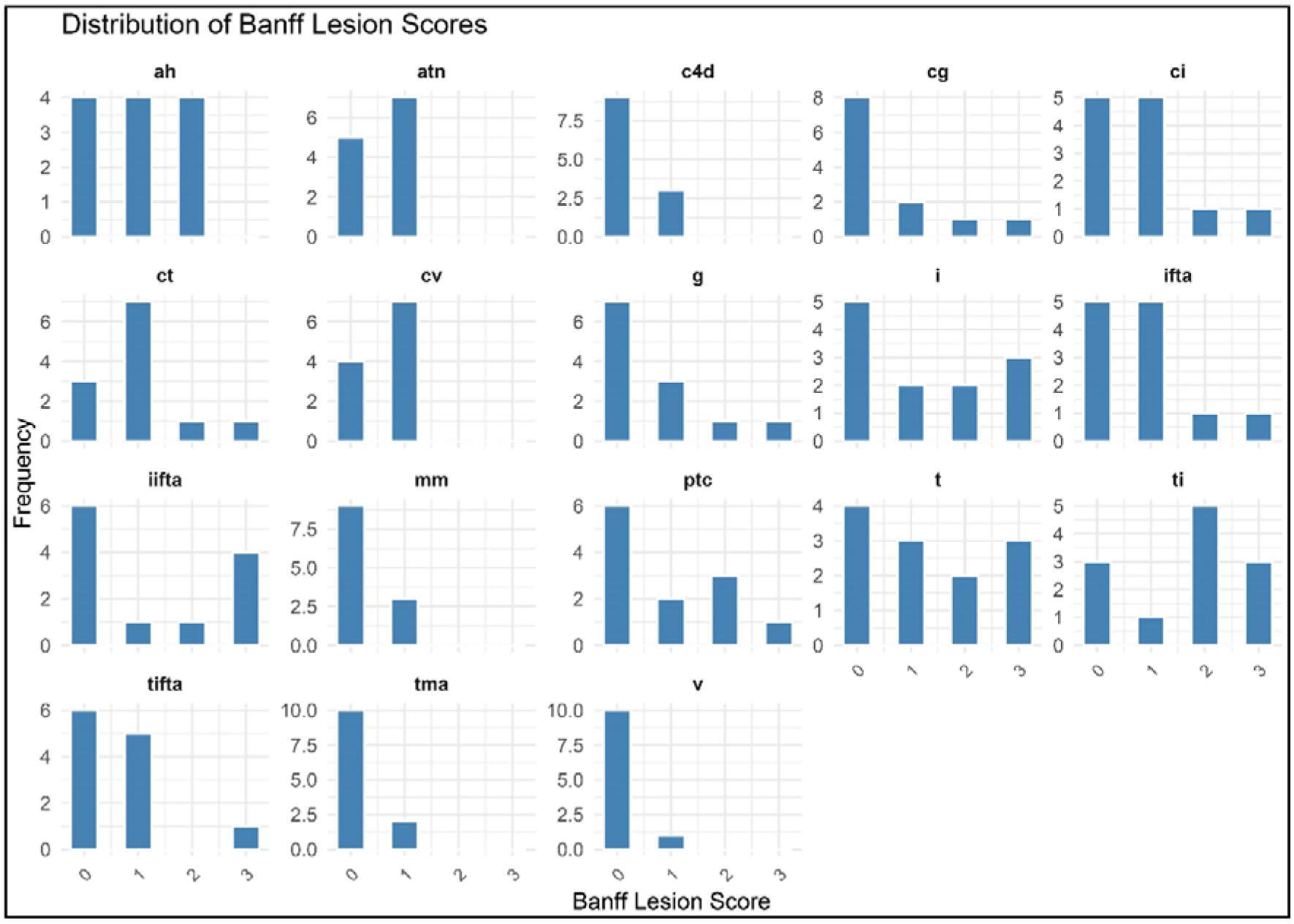
Frequency Distributions of Banff Lesion Scores. Histogram panels showing the frequency of Banff lesion scores for each lesion type across all samples.

